# Modularity and selection of nectar traits in the evolution of the selfing syndrome in *Ipomoea lacunosa* (Convolvulaceae)

**DOI:** 10.1101/2021.06.14.448377

**Authors:** Irene T. Liao, Joanna L. Rifkin, Gongyuan Cao, Mark D. Rausher

**Author notes:** Corresponding author: Irene T. Liao.

## Abstract

- Although the evolution of the selfing syndrome often involves reductions in floral size, pollen, and nectar, few studies of selfing syndrome divergence have examined nectar. We investigate whether nectar traits have evolved independently of other floral size traits in the selfing syndrome, whether nectar traits diverged due to drift or selection, and the extent to which quantitative trait locus (QTL) analyses predict genetic correlations.
- We use F5 recombinant inbred lines (RILs) generated from a cross between *Ipomoea cordatotriloba* and *I. lacunosa*. We calculate genetic correlations to identify evolutionary modules, test whether traits have been under selection, identify QTLs, and perform correlation analyses to evaluate how well QTL properties reflect the genetic correlations.
- Nectar and floral size traits form separate genetic clusters. Directional selection has acted to reduce nectar traits in the selfing *I. lacunosa*. Calculations from QTL properties are consistent with observed genetic correlations.
- Floral trait divergence during mating system syndrome evolution reflects independent evolution of at least two evolutionary modules: nectar and floral size traits. This independence implies that adaptive change in these modules requires direct selection on both floral size and nectar traits. Our study also supports the expected mechanistic link between QTL properties and genetic correlations.

## INTRODUCTION

Phenotypic divergence often involves changes in multiple suites of characters. Characters within a suite are often developmentally or genetically integrated and form an evolutionary module, and these characters evolve in a coordinated fashion (Wagner & Altenberg, 1996; Brandon, 1999; Armbruster *et al.*, 2014). By contrast, divergence of different modules can be more independent. In quantitative genetic terms, traits within modules evolve in a coordinated fashion because they are constrained by strong genetic correlations (Lande and Arnold, 1983; Cheverud, 1984; Arnold, 1992; Armbruster *et al.*, 2014), whereas different modules evolve independently because genetic correlations between modules are weak (Lewontin, 1978; Brandon, 1999). In particular, adaptive change in different modules requires selection to act directly on traits of those modules.

The degree to which complex characters are composed of distinct evolutionary modules remains an important question in evolutionary biology. In plants, floral traits and vegetative traits generally constitute distinct evolutionary modules (Berg, 1960; Armbruster *et al.*, 1999; Juenger *et al.*, 2005; Ashman & Majetic, 2006; Conner *et al.*, 2014). Flowers themselves are complex structures that serve multiple functions, from attracting and rewarding pollinators to promoting efficient pollen-to-ovule transfer for reproduction (Armbruster *et al.*, 2004; Ordano *et al.*, 2008; Diggle, 2014). However, there is much less evidence indicating whether different types of floral traits, such as flower size, nectar production, and pollen production, constitute distinct evolutionary modules.

Whether flowers consist of distinct evolutionary modules is of particular importance for understanding the evolution of pollination and mating system syndromes, which typically involves predictable shifts in floral size and shape and nectar and pollen production (Smith, 2016; Wessinger & Hileman, 2016). This predictability could arise for either of two reasons: (1) all floral traits are highly correlated genetically; or (2) flowers consist of distinct evolutionary modules that undergo similar patterns of selection in different lineages. The primary objective of this study is to distinguish between these explanations by asking whether nectar traits constitute an evolutionary module separate from other floral traits in the evolution of the selfing syndrome.

Floral-size traits and nectar traits may not constitute distinct modules given that multiple studies have demonstrated across-species correlations between aspects of flower size and nectar production (Galetto & Bernardello, 2004; Stuurman *et al.*, 2004; Kaczorowski *et al.*, 2005; Galliot *et al.*, 2006; Katzer *et al.*, 2019). One two-part hypothesis that could explain this correlation is that (1) both nectar volume and total sugar content (the product of nectar sugar concentration and nectar volume) are proportional to nectary size; and (2) genetic changes that cause smaller flowers also produce smaller nectaries by broadly affecting the cell size or cell number for all floral tissues. If both criteria are true, we expect that nectar traits would not constitute a separate evolutionary module distinct from floral size traits. However, if either criterion is false, then nectar traits may evolve independently of floral-size traits. Because nectary size has rarely been included in studies of trait divergence in pollination syndromes (but see Stuurman *et al.*, 2004; Katzer *et al.*, 2019; Edwards *et al.*, 2021), this hypothesis has not been evaluated for the evolution of any syndrome.

The repeated evolution of the selfing syndrome (Barrett, 2002; Arunkumar *et al.*, 2015) has fostered studies of its evolution in a wide range of flowering species, including *Capsella* (Slotte *et al.*, 2012; Sas *et al.*, 2016; Fujikura *et al.*, 2017; Wozniak *et al.*, 2020), *Collinsia* (Baldwin *et al.*, 2011; Strandh *et al.*, 2017; Frazee *et al.*, 2021), *Mimulus* (Fenster & Ritland, 1994; Fishman *et al.*, 2002, 2015), and *Ipomoea* (Duncan & Rausher, 2013b; Rifkin *et al.*, 2019b, 2021). Although several studies have identified quantitative trait loci (QTLs) to assess the degree to which the evolution of different syndrome traits have evolved in a correlated fashion, most have focused on herkogamy, floral size, time to flowering, and pollen size and number (Rifkin *et al.*, 2021). Because only one examined nectar production (nectar volume; Rifkin *et al.*, 2021), the extent to which nectar traits constitute a distinct evolutionary module remains to be examined. In this study, we address this issue for a pair of morning glory species, *Ipomoea cordatotriloba* and its highly selfing sister species *I. lacunosa*, which exhibits the selfing syndrome.

A second issue that we address is whether reduced nectar production in the selfing *I. lacunosa* was caused by selection acting directly or indirectly through correlated traits. Rifkin *et al.* (2019b) demonstrated that selection was responsible for reduced nectar production in this species but could not differentiate between direct or indirect selection on this trait. A finding that nectar traits constitute a distinct evolutionary module would imply that much of that selection acted directly on nectar traits. Conversely, if nectar and floral dimension traits constitute one evolutionary module, much of the selection to reduce nectar production may have been a correlated response to selection to reduce floral size.

A third issue we address is the extent to which genetic architecture can be inferred from QTL analysis. Numerous studies have identified QTLs for floral traits that differ between two species, and many have made inferences about the genetic architecture of divergence based on QTL overlaps (Slotte *et al.*, 2012; Wessinger *et al.*, 2014; Kostyun *et al.*, 2019). However, studies rarely evaluate the validity of these inferences. One exception is Gardner and Latta (2007), whose analysis was restricted largely to crop species and within-species variation.

To address these issues, we ask the following specific questions: (1) Do nectar traits constitute a separate evolutionary module from floral size traits? (2) Do correlations among nectar traits support the hypothesis that nectar volume and sugar content change primarily because of nectary size? (3) Did direct selection act to reduce nectar traits in *I. lacunosa*? and (4) How well do QTL properties predict the genetic architecture of divergence?

## MATERIALS AND METHODS

### Study System

*Ipomoea lacunosa* L. and *Ipomoea cordatotriloba* Denn. (Convolvulaceae) are sister species in series *Batatas* (Muñoz-Rodríguez *et al.*, 2018). Both are weedy species and grow widely in southeastern United States (Duncan & Rausher, 2013a; Rifkin *et al.*, 2019a; USDA, NRCS, 2021). *Ipomoea lacunosa* is highly selfing (Duncan & Rausher, 2013b), and compared to *I. cordatotriloba*, displays components of the selfing syndrome, including reductions in overall flower size, pollen amount, pigmentation, and nectar production (McDonald *et al.*, 2011; Rifkin *et al.*, 2019b; Fig. 1a).

**Fig. 0.1.**
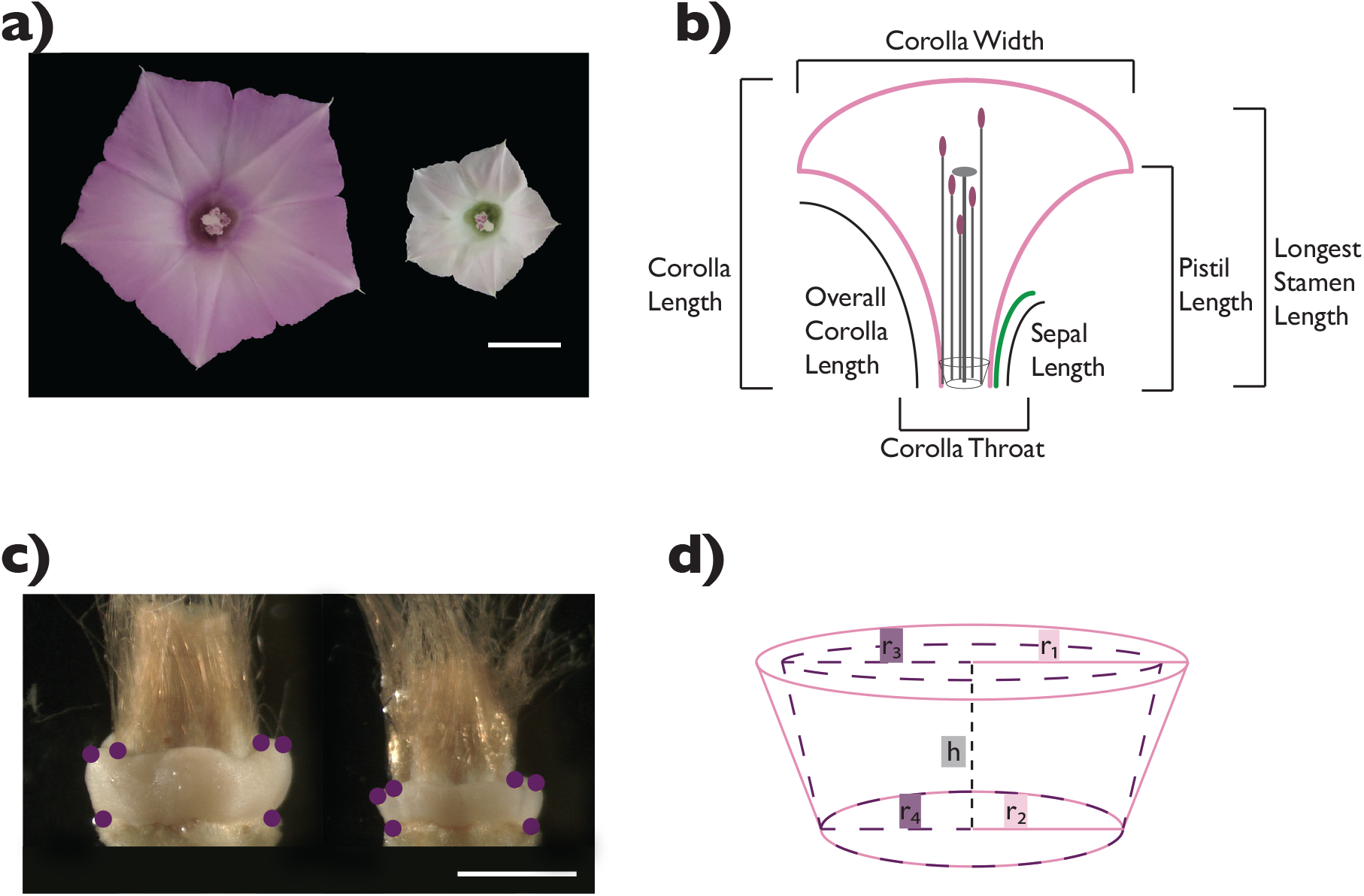
Flowers and traits measured in the study. a) Images of the two sister morning glory species used in the study, *Ipomoea cordatotriloba* (left) and *I. lacunosa* (right). Scale bar = 1 cm. b) Floral dimension traits measured in the study. c) Image of the nectary (cream-colored tissue surrounding the ovary) with purple points as landmarks for estimating the nectary size. *Ipomoea cordatotriloba* (left) and *I. lacunosa* (right). Scale bar = 1 cm. d) Nectary size approximated as the volume of the outer frustrum minus the inner frustrum. Formula for calculating the nectary size is as follows: 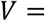 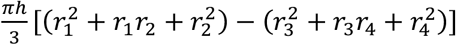, where *h* is the nectary height, *r_1_* is the nectary top outer radius, *r_2_* is the nectary top inner radius, *r_3_* is the nectary bottom outer radius, and *r_4_* is the nectary bottom inner radius.

### Materials and growing conditions

One *I. cordatotriloba* individual was crossed by one *I. lacunosa* individual to generate F1 hybrids (Duncan & Rausher, 2013b; Rifkin *et al.*, 2021). One F1 (CL5) was selfed to generate a mapping population of 500 F2s; these were selfed by single seed decent for 3 more generations to generate 322 recombinant inbred lines (RILs) at the F5 generation. Two individuals per RIL and 24-25 selfed offspring of each parent were selected for genotyping and phenotyping.

Scarified seeds were planted directly into soil in 24-cell seed packs in randomized positions and grown under 12-hour light:12-hour dark at 21-23°C to induce floral bud formation. After a month, the plants were transferred to the Duke University Greenhouses and grown under the following conditions: 12-hour light:12-hour dark, light temperatures 23-26°C, dark temperatures 16-19°C.

### F5 Phenotyping

Prior to planting, three seed traits — length, width, and mass — were measured using calipers and a balance sensitive to 0.001 grams. Seven flower-size traits were measured on open flowers using a digital caliper (“Mitutoyo Digimatic CD6”CS): corolla width, corolla throat, corolla length, overall corolla length, sepal length, longest stamen length, pistil length (Fig. 1b). Three nectar-related traits were measured: nectar volume, nectar sugar concentration, and nectary size. For 675 out of 686 total individuals, floral and nectar traits were measured on at least three flowers per individual plant.

Methods for measuring nectar volume and sugar concentration are detailed in Rifkin *et al.* (2019a) (Methods S1). Flower buds were capped the day before flower opening to limit nectar evaporation. Nectar was collected using 2μl microcapillary tubes (Drummond Scientific), and the height of the liquid in the tubes was measured with a caliper to determine nectar volume. All nectar was expelled onto a Master-53M ATAGO refractometer to measure sugar concentration. Nectaries were photographed and images were measured by selecting six points to designate the outline of the nectary (Fig. 1c). The distance between sets of points was calculated to obtain the variables to estimate the nectary size, which is modeled after the volume of a frustrum (Fig. 1d).

### Genotyping and sequencing data

Most phenotyped individuals were also genotyped (Table S1). DNA was extracted using the GeneJET ThermoFisher Plant Genomic DNA Purification Kit (Thermo Fisher Scientific). We used a modified double-digest restriction assisted DNA (ddRAD) library preparation protocol (Peterson *et al.*, 2012) combined with the Rieseberg lab genotype-by-sequencing protocol (Ostevik, 2016; Methods S2). All samples were pooled in equal amounts (100 ng) before sequencing over 4 lanes of the Illumina HiSeq 4000 platform (150bp PE reads) at the Duke Center for Genomic and Computational Biology Sequencing and Genomic Technologies Core. Raw sequence data are deposited in the NCBI Sequence Read Archive [accession: PRJNA732507].

### Sequence processing and genetic map

Markers determined from ddRAD sequencing reads were aligned to the draft *I. lacunosa* genome using NextGenMap (Sedlazeck *et al.*, 2013), and a linkage map for F5 individuals was constructed concurrently with F2 individuals (Rifkin *et al.,* 2021) using Lep-Map3 (Rastas, 2017; Methods S3). This process resulted in a total of 6056 markers with 174-625 markers per linkage group (Supporting Information Fig. S1). A “consensus” genotype was found at each position for each RIL by comparing the genotypes between the two individuals; if the genotypes were the same, the genotype was kept. All other scenarios were coded as missing data.

Overall, there was 11% missing genotype data.

### Summary statistics

The final dataset includes a total of 635 individuals phenotyped: 313 lines where both replicates were phenotyped; 9 lines where only one individual was phenotyped; 624 individuals where at least three flowers were phenotyped and 11 individuals where only one or two flowers were measured. Summary statistics were calculated in R version 4.0.2 (R Core Team, 2020).

### Correlations & heritabilities

Genetic correlations were calculated for all traits on F5 RILs in two ways: (1) as the correlation of line means, and (2) from variance and covariance components in a Multivariate Analysis of Variance, in which line is the main effect (Falconer and Mackay, 1996; Methods S4). Because these correlations are calculated from inbred lines, they represent broad-sense genetic correlations. Broad-sense heritabilities were calculated for each trait in a similar fashion (Methods S4).

### Cluster analysis

We used cluster analysis to identify evolutionary modules using three algorithms: Complete, Ward, and McQuitty with the function *hclust* (Müllner, 2013) in R. All gave similar results (Fig. S2). We computed the average and standard error of pairwise trait genetic correlations within and between each module. As an indication of whether the identified modules are real, we performed a permutation test (Methods S5).

### QTL analysis

QTL analyses were performed using both *qtl* (Broman *et al.*, 2003) and *qtl2* (Broman *et al.*, 2019) with the consensus genotypes and the average phenotypic values for each RIL. Most traits were approximately normally distributed, except for seed mass, seed width, nectar volume, and nectary size, which have slightly skewed distributions (Fig. S3).

We first used *qtl2* to identify QTLs using the scan1 function with a linear mixed model leave-one-chromosome-out (LOCO) method (Yang *et al.*, 2014; Broman *et al.*, 2019) with genome-wide LOD significance thresholds (⍺= 0.05, 1,000 permutations) and chromosome-wide LOD significance thresholds (⍺= 0.05, 10,000 permutations). We then used a multiple QTL mapping approach in *qtl* with the Haley-Knott regression method to determine non-spurious QTLs using the scantwo function to determine the penalties at ⍺=0.05 to use in the stepwiseqtl function. We started the multiple QTL model selection with genome-wide significant QTLs identified with the LOCO method, searching for only for additive QTLs. Confidence intervals were estimated at 1.5-LOD intervals.

We created two QTL datasets from these approaches. The first (designated “GWS QTL”) combined the QTLs significant genome-wide from the LOCO method and from the multiple QTL model. The second dataset (“ALL QTLs”) included all QTLs significant at the chromosome-wide thresholds, including multiple QTL peaks within a chromosome and most GWS QTLs, from the LOCO method.

We calculated the relative homozygous effect (RHE) as the difference between the mean trait values of homozygotes at the trait QTL peak divided by the difference between the mean trait values of the two original parents. Summing the RHE values for a trait provides an indication of the completeness of QTL discovery: A value substantially less than 1.0 indicates that a substantial number of QTLs have not been detected, whereas a value near 1.0 suggests most QTLs affecting a trait have been identified.

Downstream analyses were performed separately for both the GWS QTL and ALL QTL datasets. We evaluate which dataset provides a better explanation of genetic architecture by assessing how well they explain genetic correlation patterns.

### Predicting genetic architecture from QTL properties

Genetic correlations provide the best estimate of genetic architecture because they include the effects of all QTLs influencing a trait. Using properties of detected QTLs to infer genetic architecture is likely to be less reliable because typically not all QTLs affecting a trait are identified in QTL studies. To determine how well the genetic architecture of divergence is predicted by QTL properties, we examined properties of QTL co-localization. We considered QTLs for two traits to co-localize if their 1.5-LOD confidence intervals overlapped (Slotte *et al.*, 2012; Wessinger *et al.*, 2014; Kostyun *et al.*, 2019). To determine whether the overall degree of overlap was greater than expected by chance, we performed a randomization test (Methods S6). We quantified the degree of QTL overlap between two traits, examined whether QTL overlap could explain the observed modularity, and determined whether there was greater overlap within modules than between modules using a permutation test. We also assessed how well QTL overlap explained the observed genetic correlations (Methods S7).

We calculated a “predicted” genetic correlation from QTL properties, *r_Q_*, using a modification of the approach presented in Gardner and Latta (2007) (Methods S8). We assessed how well *r_Q_* matched the estimated true genetic correlations (*r_G_*) by the correlation between *r_Q_* and *r_G_*. We also calculated the correlation between average total RHE for a trait pair and bias (*r_Q_* − *r_G_*) (Gardner and Latta, 2007). To determine the significance of these correlations, we performed a permutation analysis (Methods S8). Finally, we determined whether bias involving floral and nectar traits differed for the two QTL sets by bootstrapping (1,000 replicates).

### Analyses of selection

To test for whether selection contributed to the divergence of the 13 traits, we used the *v* test from Fraser, 2020 (Methods S9) and the QTL-EE sign test (Orr, 1998). Because the QTL-EE test cannot detect selection if there are fewer than eight QTLs for a trait (Fraser, 2020), we applied this test only to traits with eight or more QTLs. Additionally, we performed simulations to evaluate the extent to which ascertainment bias might contribute to an increased false discovery rate (Results and Methods S1, Anderson & Slatkin, 2003).

## RESULTS

### Phenotypic differences

Trait differences between parent individuals, as measured on their selfed offspring, were consistent with a previous study (Rifkin *et al.*, 2019b) showing that *I. lacunosa* and *I. cordatotriloba* differ in multiple flower size and nectar traits (all traits differ significantly, P < 0.05 after correcting for multiple hypotheses (Benjamini & Hochberg, 1995; Table 1). Of the traits not examined previously, nectary size was smaller, and seed mass, seed width, and seed length were larger in the selfing *I. lacunosa* compared to the mixed-mating *I. cordatotriloba*.

**Table 0.1.** Summary of means and standard errors of 13 traits. Means and standard errors were calculated for *Ipomoea cordatotriloba* (n=25; 24 selfed offspring, 1 parent) and *I. lacunosa* (n=26; 25 selfed offspring, 1 parent) and F5 RILs (F5, n = 322). All traits are significantly different (P < 0.05) from a t-test and corrected for multiple hypotheses testing (Benjamini & Hochberg, 1995). Asterisks indicate the traits (all seeds) that do not include the parents in the mean and standard error calculations.

| Category | Trait | Units | *Ipomoea cordatotriloba* | F5 hybrids | *Ipomoea lacunosa* |
| --- | --- | --- | --- | --- | --- |
| Seed | Seed Length* | mm | 4.194 (0.0378) | 4.416 (0.0182) | 4.621 (0.036) |
| Seed | Seed Mass* | g | 0.0242<br>(0.000678) | 0.0285<br>(0.000236) | 0.0292<br>(0.000555) |
| Seed | Seed Width* | mm | 3.836 (0.0383) | 4.016 (0.0142) | 4.156 (0.0322) |
| Nectar | Nectar Sugar Concentration | mg/mL | 382.845 (2.729) | 311.498 (1.75) | 247.743 (6.474) |
| Nectar | Nectar Volume | μl | 3.239 (0.131) | 1.55 (0.0301) | 0.517 (0.0445) |
| Nectar | Nectary Size | mm<sup>3</sup> | 0.186 (0.0077) | 0.102 (0.0016) | 0.058 (0.00251) |
| Flower size | Sepal Length | mm | 11.539 (0.0965) | 11.085 (0.0432) | 10.876 (0.137) |
| Flower size | Corolla Width | mm | 28.101 (0.288) | 21.858 (0.107) | 16.424 (0.263) |
| Flower size | Corolla Throat | mm | 7.446 (0.0769) | 6.509 (0.0267) | 5.588 (0.0667) |
| Flower size | Pistil Length | mm | 16.025 (0.163) | 13.945 (0.0758) | 10.748 (0.145) |
| Flower size | Longest Stamen Length | mm | 19.418 (0.111) | 15.954 (0.0653) | 12.467 (0.114) |
| Flower size | Corolla Length | mm | 26.606 (0.198) | 22.566 (0.0906) | 17.526 (0.214) |
| Flower size | Overall Corolla Length | mm | 32.357 (0.244) | 26.573 (0.106) | 20.066 (0.277) |

### Nectar traits

Nectar volume (NV) and total sugar amount (TS) are highly correlated (*r* = 0.98; Fig. S4b,c). Both nectar volume and total sugar amount also exhibit significantly non-linear relationships with nectary size, with both regressions having high R^2^ values (~0.92), indicating that the regressions account for much of the variation seen in nectar volume and total sugar amount (Fig. S4e,f). From these regressions, we derived a relationship that predicted sugar concentration (NSC) as a function of nectary size (NS): NSC(NS) = TS(NS)/NV(NS). This predicted relationship is very similar to the relationship between measured sugar concentration and nectary size (Fig. S4g). Because total sugar amount is a derived value with a nearly exact correspondence with nectar volume, we do not include total sugar amount in subsequent analyses.

### Genetic correlations and evolutionary modules

Genetic correlations calculated from the variance-covariance components are similar to those calculated from the RIL means (*r* = 0.981, p < 0.05, Fig. S5). We thus used only the former in subsequent analyses.

Pairwise genetic correlations are positive for all traits, as are most pairwise environmental correlations (Table S2, Fig. S6b,c). The three clustering algorithms produced similar results: The 13 traits form three distinct clusters, which we interpret as evolutionary modules. The “flower module” consists of all floral size traits, the “nectar module” consists of the three nectar traits, and the “seed module” consists of the three seed-size traits (Fig. 2, S2). The average correlations within each module are substantially higher (range 0.557-0.656) than the average correlations between traits in different modules (range 0.008 — 0.226; Table 2a). The average within-module correlation (0.602) is more than 5 times the average between-module correlation (0.117) (Table 2a).

**Fig. 2.**
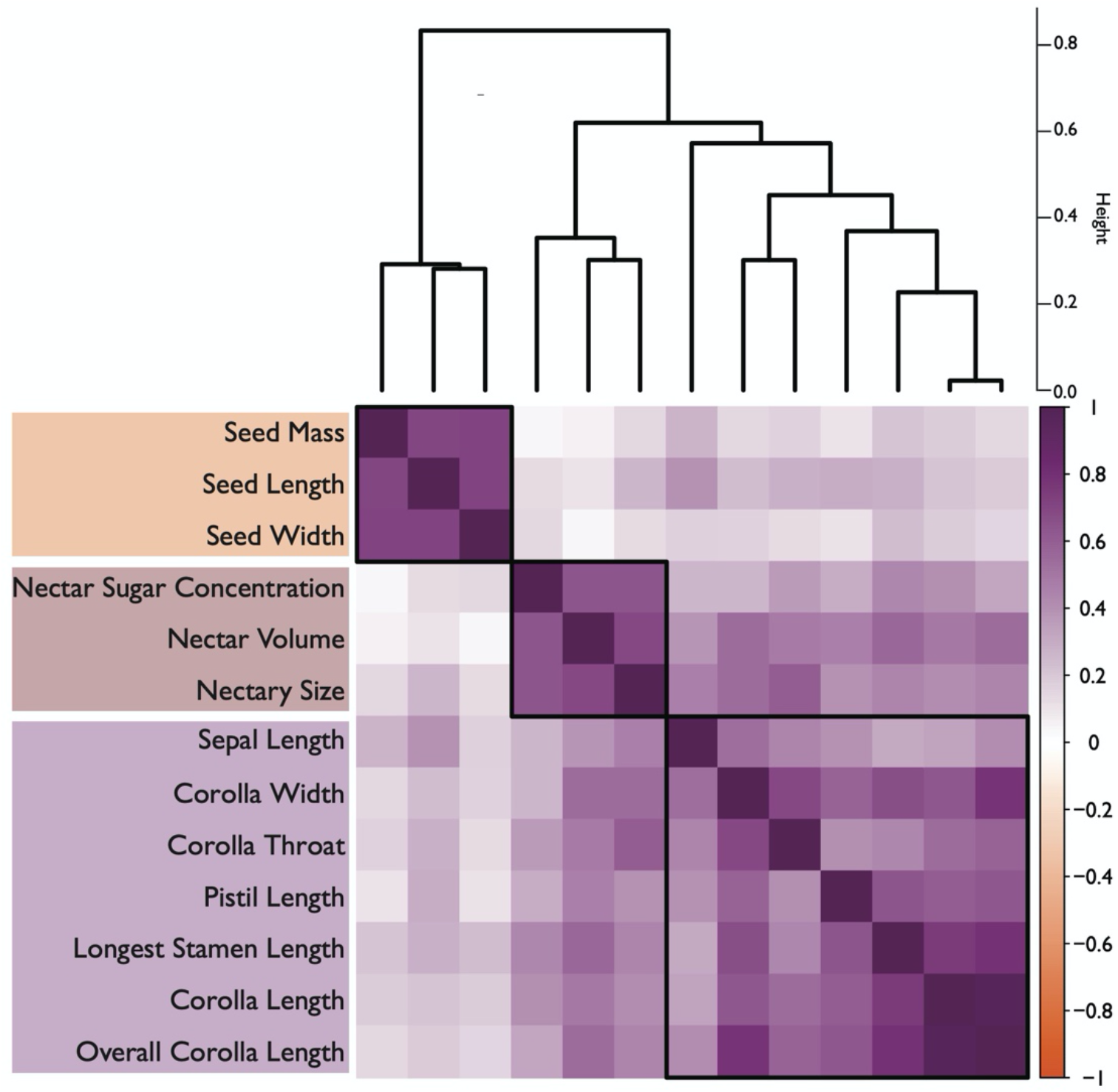
Cluster diagram and heatmap based on variance-covariance-component genetic correlations. Cluster dendrogram portrayed for the “mcquitty” method. Boxes in the heat map indicate the three clusters. Other clustering methods result in the same three clusters (Fig. S6). For the heatmap, strong positive correlations are in dark purple while weak to no correlation range from light purple to white. (left to right) seed, nectar, and flower size. Pairwise genetic correlation values are found in Table S1.

**Table 0.2.** Average genetic correlations and QTL overlap within and between modules. a) Average genetic correlations calculated from variance and covariance components. Average within-and between-module correlations are 0.602 and 0.117, respectively. Permutation test of hypothesis that within-module correlations do not differ from between-module correlations: P < 0.001. b) Average QTL overlap for GWS QTLs. Average within-and between-module correlations are 0.248 and 0.086, respectively. Permutation test of hypothesis that within-module correlations do not differ from between-module correlations: P = 0.078. c) Average QTL overlap for ALL QTLs. Average within-and between-module correlations are 0.464 and 0.275, respectively. Permutation test of hypothesis that within-module correlations do not differ from between-module correlations: P = 0.047.

**a) GENETIC CORRELATIONS**
| MODULE | seed | nectar | flower |
| --- | --- | --- | --- |
| seed | 0.651 (0.0655) | 0.117 (0.0224) | 0.0077 (0.0087) |
| nectar | 0.117 (0.0224) | 0.557 (0.1189) | 0.226 (0.0177) |
| flower | 0.0077 (0.0087) | 0.226 (0.0177) | 0.601 (0.0381) |

**b) QTL OVERLAP – GWS QTLs**
| MODULE | seed | nectar | flower |
| --- | --- | --- | --- |
| seed | 0.111 (0.111) | 0.041 (0.023) | 0.080 (0.015) |
| nectar | 0.041 (0.023) | 0.112 (0.082) | 0.110 (0.014) |
| flower | 0.080 (0.015) | 0.110 (0.014) | 0.286 (0.031) |

**c) QTL OVERLAP – ALL QTLs**
| MODULE | seed | nectar | flower |
| --- | --- | --- | --- |
| seed | 0.460 (0.111) | 0.216 (0.049) | 0.214 (0.031) |
| nectar | 0.216 (0.049) | 0.500 (0.096) | 0.360 (0.024) |
| flower | 0.214 (0.031) | 0.360 (0.024) | 0.459 (0.028) |

Generally, this pattern suggests that evolution of the three modules has been largely genetically independent. Nevertheless, a moderate average genetic correlation (0.226) exists between floral and nectar traits (Table 2a), suggesting that there may have been some correlated evolution of these two modules.

A permutation test that randomly assigned the observed correlations to trait pairs indicated that the modules identified reflect groups with correlations that are higher than expected by chance. In 1,000 permutations, no values of the difference between average correlations within and between modules was as large as the observed difference of 0.485, allowing rejection of the null hypothesis at P < 0.001.

### QTL analyses: number and effect sizes

Combining QTLs with genome-wide significance from the LOCO model and the multiple QTL model (GWS QTLs), we identified 97 QTLs. From just the LOCO model with chromosome-wide significance (CWS), ALL QTLs produced total of 146 QTLs (92 GWS and 54 CWS; Fig. 3, S7, Table S3). QTLs were located throughout the genome, with 3-12 QTLs per chromosome for GWS QTLs and 4-17 QTLs for ALL QTLs. Most traits have more than 5 QTLs detected, except for seed mass and seed width (in both QTL datasets) and nectar volume and seed length (in GWS QTLs).

**Fig. 3:**
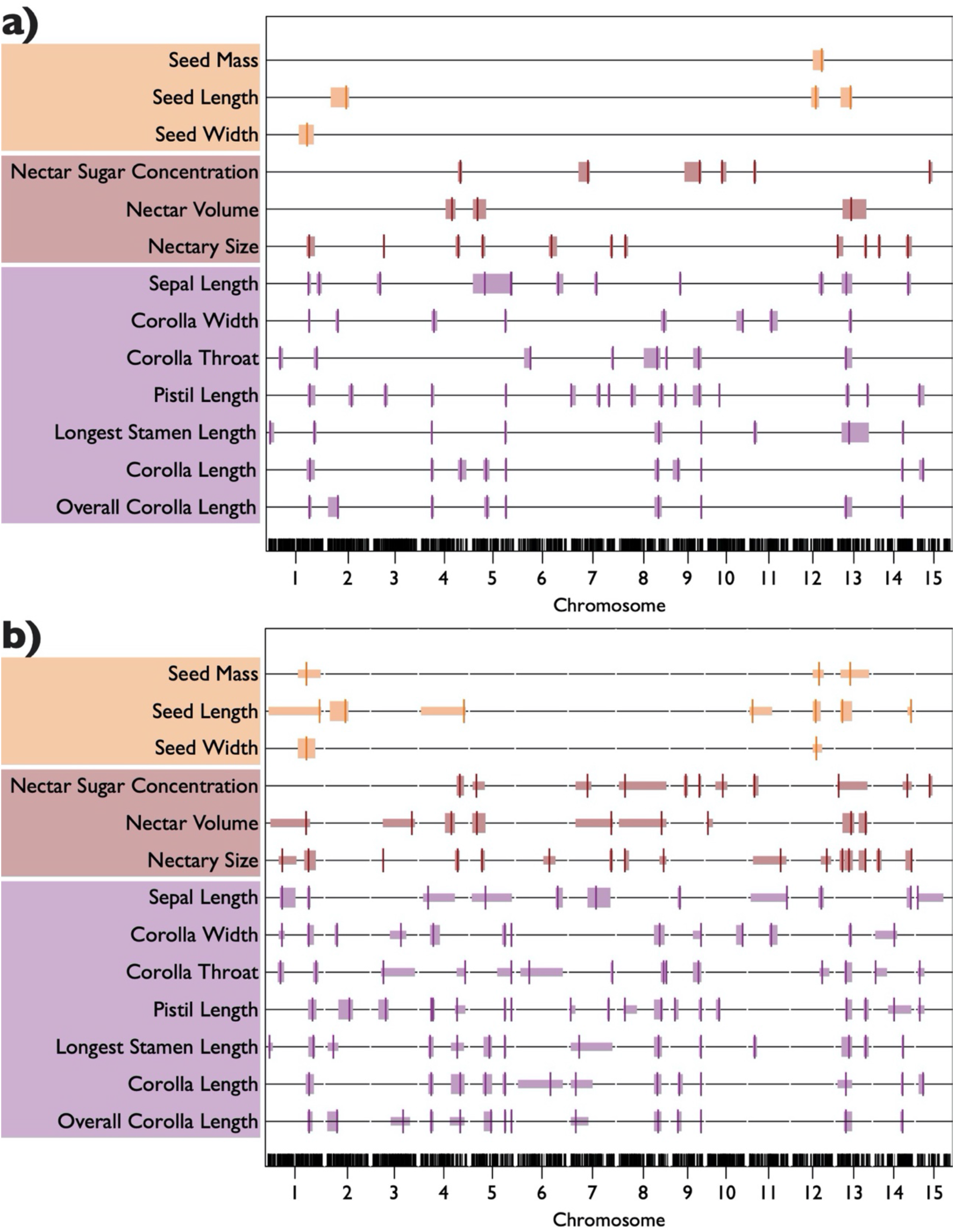
Chromosome map of QTLs for 13 phenotypic traits. Each bar represents the 1.5 LOD confidence interval with the vertical line indicating the QTL peak. Bars in orange hues represent seed traits; maroon red hues represent nectar traits; purple hues represent flower traits. A summary of QTL peaks is found in Table S2. Individual trait QTL plots are found in Fig. S7. a) Genome-wide significant (GWS) QTLs – combination of the two most stringent QTL models. b) ALL QTLs – based on multiple QTL peaks from the LOCO method. Thicker bars indicate peaks significant at the genome-wide threshold; thinner bars indicate peaks significant only at the chromosome-wide threshold.

Most QTLs are of small to moderate effect (Table S3, S4). For GWS QTLs, mean RHE for most traits was less than 0.15, except for seed mass (0.375) and seed width (−0.372). The maximum RHE was less than 0.25 for nine traits and was less than 0.5 for the three seed traits. Only sepal length had a large-effect QTL (RHE = 0.891). The pattern was similar for ALL QTLs: all traits had a mean RHE less than 0.15 in absolute value, nine traits had a maximum RHE less than 0.25, and seed traits had a maximum RHE less than 0.5.

For floral traits, total RHE values are relatively high (GWS QTLs: 0.461 − 1.134, mean = 0.802; ALL QTLs: 0.721 − 1.562, mean = 1.013), suggesting that the identified QTLs account for much of the difference between the two species (Table S3b). Less is accounted for by nectar traits (GWS QTLs: 0.247 − 0.654, mean = 0.436; ALL QTLs: 0.563 − 0.904, mean = 0.740), indicating the existence of an unknown number of undetected QTLs. Finally, we were least successful in recovering QTLs for seed traits (GWS QTLs: −0.372 − 0.375, mean = 0.078; ALL QTLs: less than 0).

### QTL overlap and genetic correlations

The pattern of average QTL overlap is similar to that exhibited by genetic correlations (Table 2). For both QTL sets, the average within-module QTL overlap was greater than that between modules (Table 2b,c). However, comparisons with genetic correlations differ somewhat for the two datasets. For GWS QTLs, the within-module QTL overlap averages are lower than, while the between-module QTL averages are similar to, the corresponding genetic correlation averages (Table 2a,b). By contrast, for ALL QTLs, the within-module overlap averages are more similar to the corresponding genetic correlation averages (Table 2a,c), while the between-module averages are somewhat higher. The within- and between-module averages of the predicted genetic correlations, *r_Q_*, also show similar patterns when compared to the actual genetic correlations (Table 2, S6).

The randomization test, in which we assigned each QTL to a random position in the genome and calculated the resulting QTL overlap (Table S7), indicated that there is greater overlap among QTLs within modules than would be expected by chance. For GWS QTLs, this test was significant (P < 0.0001) across all modules combined. However, this result is likely primarily due to overlap found within the floral-size module (P < 0.0001; Table S7a); neither the nectar nor the seed modules differ from the null hypothesis of random placement of QTLs in the genome (Table S7a). By contrast, for ALL QTLs the test is significant both over all the modules combined and for each module individually (Table S7b). These results are consistent with within-module pleiotropy of QTLs.

For individual trait pairs, rather than the module averages considered above, QTL overlap and the predicted genetic correlations, *r_Q_*, are generally reflective of the magnitude of the corresponding genetic correlations, *r_G_*, for both GWS and ALL QTLs. The correlation between *r_G_* and the QTL overlap is significant for both GWS QTLs and ALL QTLs (Fig. 4), as is the correlation between *r_G_* and *r_Q_* (Fig. 5a). A subset of *r_Q_* values is negative and correspond largely to pairwise correlations that include at least one seed trait. Omitting seed traits, the correlation between *r_G_* and *r_Q_* is reduced somewhat for both QTL sets but remain highly significant (Fig. 5b). These patterns suggest that the magnitude of QTL overlap and the predicted genetic correlations capture the actual genetic correlation between traits reasonably well.

**Fig. 4.**
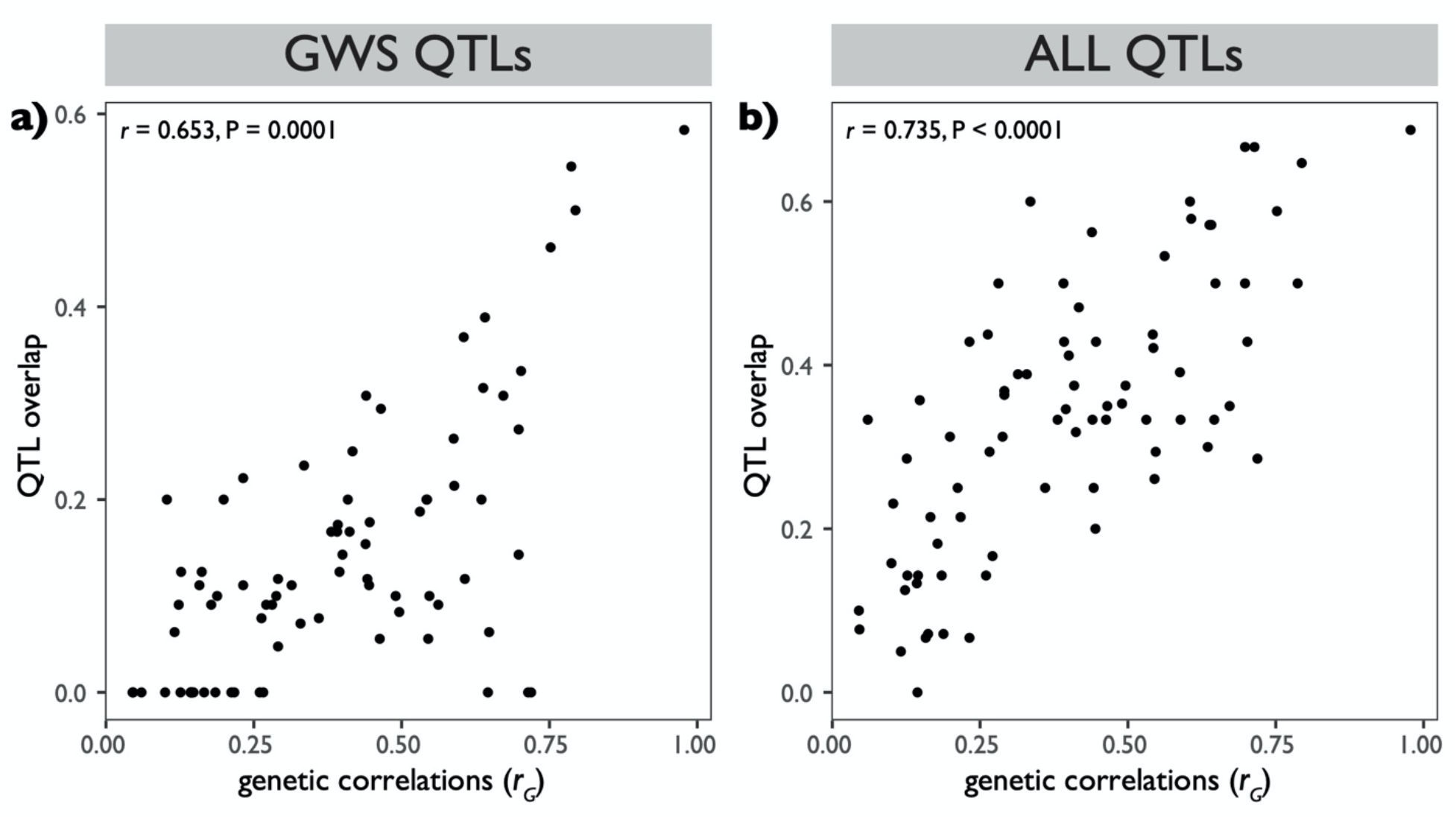
Comparison of genetic architecture based on genetic correlations versus QTL overlap. Figures portray correlation between genetic correlations and QTL overlap for all trait pairs. a) QTL overlaps based on GWS QTLs only. b) QTL overlaps based on ALL QTLs.

**Fig. 5.**
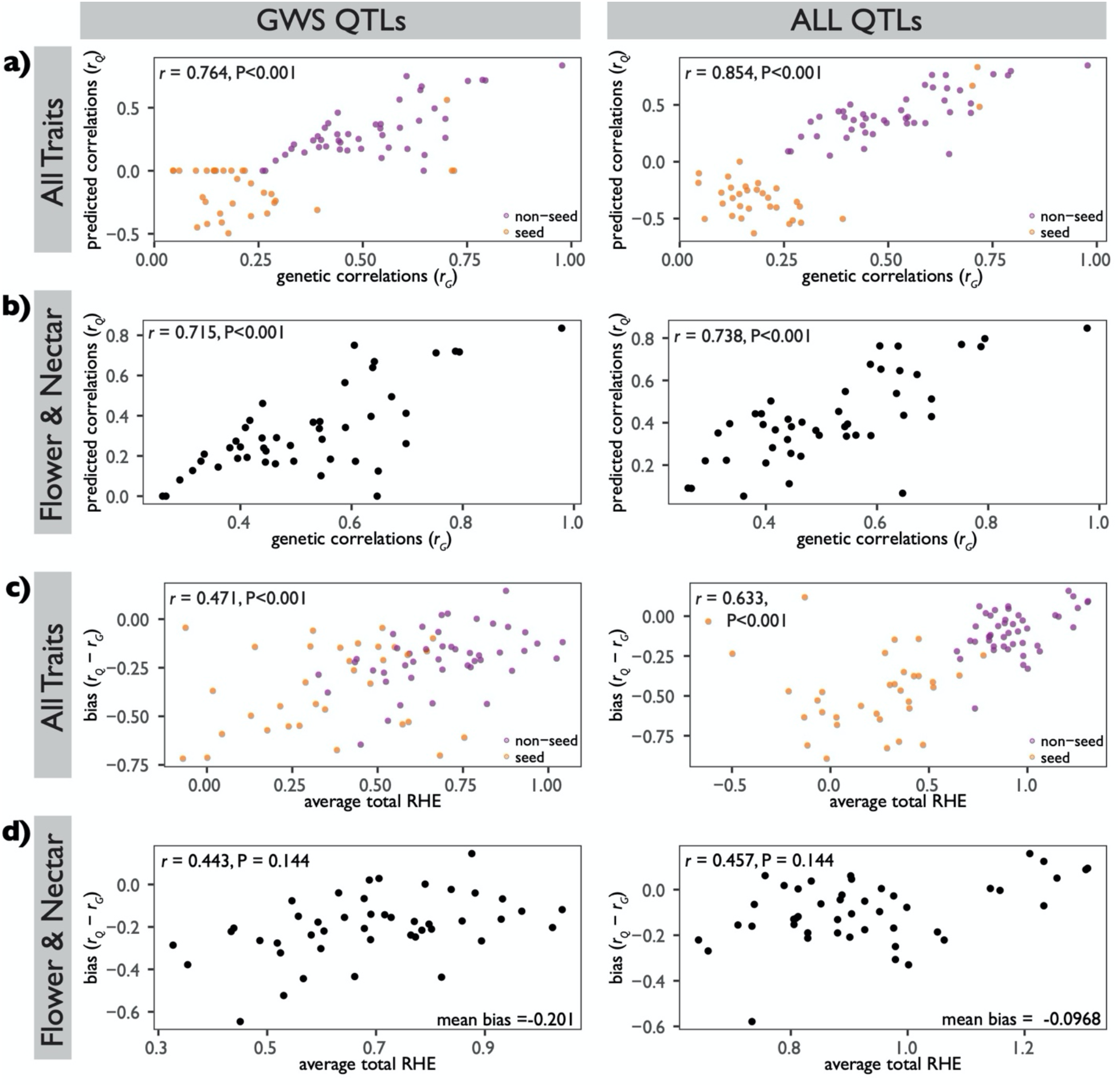
Accuracy of predicted correlations. The average total RHE was calculated by taking the average of total RHE for the two traits. Significance of correlation was determined by a permutation test. a) Correlation between *r_G_* and predicted genetic correlations from QTL effects (*r_Q_*) for all traits. b) Same as a) but excluding seed traits. c) Correlation between bias and average total RHE for all traits. d) Same as c) but excluding seed traits. For a) and c), orange points indicate correlations that include one of the seed traits; purple points indicate correlations of non-seed traits (floral and nectar traits only). For all figures, comparisons for GWS QTLs are on the left; those for ALL QTLs are on the right.

The accuracy of the predicted genetic correlation is reflected in the bias, *r_Q_* - *r_G_*. For both GWS QTLs and ALL QTLs, bias was significantly less for trait pairs with greater total RHE (Fig. 5c). Although this pattern holds after removing the correlations that include seed traits, the permutation test is not significant for either QTL set (Fig. 5d). For just nectar and floral size traits, average bias when ALL QTLs were used (−0.0968) was about half the bias when just GWS QTLs were used (−0.201), a difference that was significant with a bootstrap comparison (P < 0.001).

### Analysis of selection

A strong preponderance of QTLs with RHE in the direction of the species difference (“consistent-directional QTLs”) compared to the opposite direction (“contra-directional QTLs”) is an expected signature of selection. Overall, 81.5% of QTLs were consistent-directional, with the proportion for ALL QTLs being larger than for GSW QTLs (Table S8), a pattern consistent with selection acting on many of the traits examined.

Both the QTL-EE sign test and the Fraser test support this inference. With the former test, for GWS QTLs, most of the floral traits have a nominally to borderline significant excess of consistent-directional QTLs, with corolla width and corolla length significant at P < 0.05 after correcting for multiple comparisons (Table S4A). For ALL QTLs, all floral size and nectar traits, except for sepal length, were found to be nominally significant, with most of the floral size traits and nectar sugar concentration remaining significant after correcting for multiple comparisons (Table S4B). The Fraser *v* test statistic is highly significant for all traits (all P < 0.002; Table 3), except sepal length and all three seed traits, and remain significant at an overall level of P < 0.01 after a sequential Bonferroni correction. Overall, these results are consistent with selection having acted on all three nectar traits and all floral traits except sepal length, and agree with findings from a previous QstFst analysis (Rifkin *et al.*, 2019b).

**Table 3.** Fraser (2020) test for selection on each trait. Columns are components of Equation [2] in Fraser (2020), *v* statistic given by that equation, and probability of *v* being as large or larger than observed by chance. A *c* value of 1.0 was used in all calculations (Methods S9). All significant P values remain significant at an overall level of P < 0.01 after a sequential Bonferroni correction for multiple comparisons (indicated by asterisks).

| | Between-Species<br>Variance<br>Component | Number<br>of<br>CORD<br>parents | CORD<br>parent<br>variance | Number<br>LAC<br>parents | LAC<br>parent<br>variance | F5<br>Between-Line<br>variance<br>component | F5 Within-Line<br>variance<br>component | H<sup>2</sup> (Broad-sense<br>heritability) | F5<br>Phenotypic<br>Variance | $\nu$ (test<br>statistic) | P<br>(significance<br>of test) |
| --- | --- | --- | --- | --- | --- | --- | --- | --- | --- | --- | --- |
| Corolla Width | 66.656 | 25 | 1.759 | 26 | 1.801 | 2.450 | 2.261 | 0.520 | 4.678 | 27.339 | 0.00001526* |
| Corolla Throat | 1.687 | 25 | 0.146 | 26 | 0.116 | 0.155 | 0.137 | 0.531 | 0.290 | 10.895 | 0.001* |
| Corolla Length | 41.030 | 25 | 1.016 | 26 | 1.186 | 1.652 | 1.849 | 0.472 | 3.489 | 24.867 | 1.5259E-05* |
| Overall Corolla Length | 74.652 | 25 | 1.434 | 26 | 1.993 | 2.515 | 2.094 | 0.546 | 4.583 | 29.795 | 1.5259E-05* |
| Sepal Length | 0.203 | 25 | 0.243 | 26 | 0.490 | 0.363 | 0.446 | 0.449 | 0.806 | 0.482 | 0.490 |
| Longest Stamen Length | 23.796 | 25 | 0.255 | 26 | 0.337 | 0.911 | 0.801 | 0.532 | 1.692 | 26.400 | 1.5259E-05* |
| Pistil Length | 13.970 | 25 | 0.686 | 26 | 0.545 | 1.480 | 0.702 | 0.678 | 2.173 | 9.443 | 0.002 |
| Seed Mass | 0.000 | 24 | 0.000 | 25 | 0.000 | 0.000 | 0.000 | 0.620 | 0.000 | 0.874 | 0.350 |
| Seed Length | 0.090 | 24 | 0.034 | 25 | 0.033 | 0.088 | 0.034 | 0.724 | 0.121 | 0.989 | 0.320 |
| Seed Width | 0.050 | 24 | 0.035 | 25 | 0.026 | 0.048 | 0.034 | 0.588 | 0.081 | 0.992 | 0.320 |
| Nectar Volume | 3.557 | 25 | 0.374 | 26 | 0.052 | 0.161 | 0.242 | 0.400 | 0.402 | 22.016 | 1.5259E-05* |
| Nectar Sugar Concentration | 9056.500 | 25 | 191.533 | 26 | 1089.830 | 602.835 | 714.065 | 0.458 | 1312.340 | 14.993 | 0.0001* |
| Nectary Size | 0.008 | 25 | 0.001 | 26 | 0.000 | 0.001 | 0.000 | 0.565 | 0.001 | 12.788 | 0.0004* |

Contra-directional QTLs weaken the ability to detect selection because the alleles fixed are assumed to be due to genetic drift. Alternatively, contra-directional QTLs may have advantageous pleiotropic effects on other characters and were fixed because of those positive effects. Most floral size and nectar contra-directional QTLs overlap with at least one other floral or nectar QTL with consistent-directional effects (Table S9). This observation is consistent with many of the contra-directional QTLs being fixed by selection, which would make the role of selection in the evolution of these traits even stronger than is revealed by the Fraser and QTL-EE sign tests.

Anderson and Slatkin (2003) demonstrate that ascertainment bias can increase the rate of false positives in the QTL-EE test and likely also for the Fraser test. However, their analyses implemented an extreme version of ascertainment bias, in which the most diverged trait was chosen out of *N* candidate traits measured. Because we believe this does not reflect how organismal traits are selected for analysis (see Results and Methods S1), we performed simulations that relax this assumption (Results and Methods S1). Briefly, we assume that traits for study are chosen randomly from the *βN* candidate loci with the highest divergence. *β* is thus an index of the intensity of ascertainment bias, with low values indicating high intensity. In Anderson and Slatkin’s study, *β* = 1/*N*. We find that to correct for ascertainment bias, the probability value from the test must be multiplied by *β* to obtain the true probability. Although there is admittedly no theoretical or empirical guidance, we believe it is unrealistic to assume that traits chosen for study are among the top 0.15 or less (see Results and Methods S1 for justification). Thus, we interpret a P value of less than 0.05 · 0.15 = 0.0075 as strong evidence for selection despite any ascertainment bias. We also believe that it may be unlikely to choose traits from among the top 0.4 most diverged traits, which would mean that a P value of less than 0.05 · 0.4 = 0.02 may be consistent with selection. Based on these criteria, our analyses indicate that, except for corolla throat and pistil length, the signatures of selection we detected on floral size and nectar traits are not likely an artifact of such bias (Results and Methods S1). More generally, our analyses indicate that ascertainment bias likely does not inflate the probability of obtaining false positives in the QTL-EE and Fraser tests nearly as much as suggested by Anderson and Slatkin (2003).

## DISCUSSION

### Nectar Traits and Floral Size Traits are Separate Evolutionary Modules

Flowers are complex structures that perform a variety of functions that are affected by floral size and shape, nectar production, and pollen production. Divergence in these traits is common between closely related species due to changes in either the predominant pollinators or favored mating system. Whether this divergence is constrained by pleiotropy and genetic correlations among floral traits is a long-standing question in plant evolutionary biology (Smith, 2016; Wessinger & Hileman, 2016; Kostyun *et al.*, 2019). While some recent studies have attempted to identify the degree of genetic correlations and modularity among floral traits among species (Dellinger *et al.*, 2019; Dellinger, 2020; Reich *et al.*, 2020), we are still largely ignorant of whether flowers consist of distinct evolutionary modules and, if so, what those modules are.

The flowers of *Ipomoea lacunosa* and *I. cordatotriloba* consist of at least two distinct evolutionary modules: floral size traits and nectar traits. A previous study of these species only examined one nectar trait (nectar volume), which clustered with floral size traits (Rifkin *et al.*, 2021). By including additional nectar traits, we could distinguish these modules by the moderately high within-module genetic correlations, but low between-module correlations (Table 2a). This pattern is also reflected in the degree of QTL overlap, which is on average higher within the two modules than between them. This genetic architecture indicates that the evolution of decreased floral size in *I. lacunosa* has been largely genetically independent of the evolution of reduced nectar production and *vice versa*.

Limited information exists in other species regarding the genetic and evolutionary independence of floral size and nectar traits. Across multiple species within the same genus, floral size is phylogenetically correlated with nectar volume (Galetto & Bernardello, 2004; Kaczorowski *et al.*, 2005; Tavares *et al.*, 2016), but such patterns are uninformative about evolutionary independence. Within species, artificial selection for floral size in *Eichhornia paniculata* resulted in a correlated response in nectar volume, indicating that size and nectar volume are genetically correlated, but the magnitude of the correlation is unknown (Worley & Barrett, 2000). Genetic correlations between aspects of flower size and nectar production in *Nicotiana alata* were non-significant (Kaczorowski *et al.*, 2008), consistent with the existence of separate floral size and nectar modules. Despite these two studies, the genetic architecture of within-species variation is not necessarily indicative of the genetic architecture of divergence for two reasons: (1) genetic correlations can change under selection (Sheridan & Barker, 1974; Mitchell-Olds & Rutledge, 1986; Falconer and Mackay, 1996; Roff, 2007; Arnold *et al.*, 2008), and (2) new mutations do not necessarily reflect the correlation structure of standing genetic variation.

A few QTL studies have examined nectar trait divergence between species, but most report only phenotypic correlations between aspects of floral size and only one or two nectar traits. These studies reveal moderate to complete QTL overlap between floral size and nectar volume. For *Petunia integrifolia* and *P. axillaris*, all four nectar volume QTLs overlap with floral size QTLs with the highest phenotypic correlation between the lower subdomain of the corolla tube and nectar volume (0.58; Galliot *et al.*, 2006). For *Ipomopsis guttata* and *I. tenufolia*, two nectar volume QTLs overlap with floral size QTLs (Nakazato *et al.*, 2013). For *Penstemon amphorellae* and *P. kunthii*, nectar volume is correlated with lateral stamen length (0.742) and nectary size (0.603; Katzer *et al.*, 2019). Finally, for *Aquilegia brevistyla* and *A. canadensis,* nectar volume and sepal traits had an average phenotypic correlation of 0.53 and an average QTL overlap of 0.4, but nectary size and sepal traits had a higher average correlation (0.673) and QTL overlap (0.53; Edwards *et al.*, 2021). While these findings are not completely in line our results, none of these studies attempt to identify evolutionary modules, nor do they consider the extent to which QTL co-localization reflects actual genetic correlations among traits. It is thus unclear whether the genetic architecture of divergence in these species differs fundamentally from that we report.

One explanation for the independence of floral size and nectar modules is differences in the developmental timing of these traits. Although the development of the flower and the floral nectary are undoubtedly linked, they may differ in the timing of cell fate specification and the coordination between cell division and expansion. The nectary often develops *after* floral organs have been specified (Smyth *et al.*, 1990; Baum *et al.*, 2001; Thornburg, 2007; Jeiter *et al.*, 2017); in these *Ipomoea* species, the nectary is not visibly present in the earliest stages of flower development when the four major floral organs are easily identifiable (personal observation).

Floral integration may be diminished in selfing species (Anderson & Busch, 2006), although a meta-study suggests otherwise (Fornoni *et al.*, 2016). By examining traits in addition to floral size, our study and a parallel study (Rifkin *et al.*, 2021) support weakening floral integration in selfing species. Rifkin *et al.* (2021) demonstrated that pollen traits (pollen number and pollen size) constitute a distinct evolutionary module. Although neither study address whether pollen and nectar traits constitute distinct modules, they raise the possibility that flowers in these species consist of at least three distinct evolutionary modules.

### Structure of the Nectar Module

Galetto and Bernadello (2004) report a correlation between nectar volume and nectary size across six *Ipomoea* species. Additionally, within *Nicotiana alata*, there is a high genetic correlation (0.89) between nectar volume and nectar energy content (total sugar amount) (Kaczorowski *et al.*, 2008). These two findings suggest that (1) physiologically, nectar volume and sugar content are proportional to nectary size and that (2) these three traits may comprise a distinct evolutionary module. If the hypothesis is true, evolutionary changes in nectar volume and sugar content could primarily be correlated responses to changes in nectary size. Although this hypothesis is not supported in the divergence for *Petunia axillaris* and *P. integrifolia*, where nectar volume and nectar sugar concentration differ between the two species but the nectary size remains the same (Stuurman *et al.*, 2004), it may be true for other species, such as *Aquilegia brevistyla* and *A. canadensis* (Edwards *et al.*, 2021).

Our results are somewhat consistent with this hypothesis. Nectar volume and total sugar content are each moderately genetically correlated with nectary size (*r* ≈ 0.65 in both cases, Table S2) and highly correlated with each other (Fig. S4). Sugar concentration is also positively genetically correlated with nectary size with *r* ≈ 0.65, and both nectar volume and nectar sugar concentration have moderately high QTL overlap with nectary size (0.66 and 0.5, respectively, Table S5). However, with genetic correlations of this magnitude, correlated responses to selection on nectary size would explain less than half of the variation in nectar volume and total sugar. The rest would be accounted for by mutations that affect volume and/or total sugar but not nectary size. Even within the nectar module, there is substantial independent evolution of the component traits.

### Selection on Floral Size and Nectar Traits

Two previous QstFst studies determined that natural selection contributed to the divergence of five floral size and two nectar traits between *Ipomoea lacunosa* and *I. cordatotriloba* (Duncan & Rausher, 2013b; Rifkin *et al.*, 2019b, 2021). However, neither study was able to distinguish between selection acting directly on these characters and indirect selection due to correlations with selected characters. The QTL-EE sign test and the Fraser test presented here also reveal that divergent selection likely operated on both floral size and nectar traits. By combining these results with our information on genetic architecture, we infer that some floral size traits and some nectar traits likely diverged due to direct selection on those traits; the weak genetic correlations between the two modules would unlikely yield detectable signatures of indirect selection. Because of the high within-module genetic correlations, we cannot distinguish between direct and indirect selection for traits within the same module. Moreover, our study is unable to identify the causes of selection acting on these traits, *i.e.* whether selection in *I. lacunosa* was favored by advantages of resource re-allocation, increased rate in floral development, or decreased florivory (Sicard & Lenhard, 2011).

Ascertainment biases, in the form of choosing to study characters that are known to have diverged between species, can bias both QTL-EE sign tests and the Fraser test, artificially increasing the probability of obtaining false positives (Anderson & Slatkin, 2003). However, we believe that ascertainment biases do not account for the apparent selection on nectar and floral size traits for three reasons. First, our results are consistent with previous selection detected by a completely different method (Duncan & Rausher, 2013b; Rifkin *et al.*, 2019b). Second, nine of ten floral size and nectar traits are highly significant (P < 0.002) by the Fraser test (Table 3), and for seven of these traits, neutrality can be rejected if they were chosen for study from more than the top 15% of candidate traits in terms of divergence (Results and Methods S1). Finally, for these seven traits, the maximum number of candidate traits from which each of those traits can be drawn for rejection of neutrality is likely much higher than the actual number of candidate traits. Generally, the results of our analyses of ascertainment bias also indicates that they likely do not inflate the probability of false positives nearly as much as suggested by Anderson and Slatkin.

### Do QTL Traits Predict the Genetic Architecture of Divergence?

QTL studies often infer properties of the genetic architecture of divergence from patterns of QTL overlap (*e.g.* Slotte *et al.*, 2012; Wessinger *et al.*, 2014; Kostyun *et al.*, 2019). Seldom are these inferences evaluated by comparing the patterns of QTL overlap and the actual genetic correlations among traits. One exception is a meta-study by Gardner and Latta (2007), which used QTL properties from published studies to predict genetic correlations among traits and compared those estimates to actual genetic correlations. Because most of the studies examined quantified patterns of within-species variation, it is unclear whether these patterns can be extended to the genetic architecture of divergence between species.

Our results indicate that QTL properties can predict genetic architecture reasonably well qualitatively. As in Gardner and Latta (2007), the magnitudes of the observed genetic correlations in our study are positively correlated with QTL overlap and with predicted genetic correlations, with the strength of the correlation being similar to or higher than reported by Gardner and Latta (Fig. 5). This conclusion also extends to the modular nature of divergence: the average QTL overlap within and between modules generally mirrors the average genetic correlations within and between modules.

Bias, the difference between the predicted and observed genetic correlation, increased as total RHE decreased. Despite the tendency for QTL effects to be overestimated in studies with fewer than 500 individuals (Beavis *et al.*, 1994; Xu, 2003), RHE can be interpreted as an index of the completeness of QTL identification. Based on this, our results indicate that the accuracy of the predicted genetic architecture is proportional to the degree to which all QTLs affecting the traits have been identified. In our study, total RHE was generally high for flower size and nectar traits, and the corresponding genetic correlations had little bias. By contrast, total RHE was low for seed traits, and the bias in the predicted correlation was substantial. We conclude that it is dangerous to make inferences about genetic architecture and constraints on evolution based on QTL overlap unless most of the QTLs affecting the traits of interest have been identified. Thus, while QTL studies can confirm the molecular underpinnings of genetic correlations, assessment of the genetic architecture of genetic correlations can be done more reliably by estimating those correlations directly.

Finally, although QTLs identified as significant only at the chromosome-wide level (CWS) are often considered less reliable than those significant genome-wide, our results indicated that the CWS QTLs we detected are likely real because including these QTLs in our analyses provided improved estimates of the genetic architecture of divergence. If CWS QTLs were artifactual, we would expect (1) correlations to *decrease*, (2) bias to *increase*, and (3) proportion of QTLs in the same direction of the species difference to be 0.5. We do not observe any of these expectations. First, the correlation between the observed genetic correlations (*r_Q_*) and QTL overlaps are higher when CWS QTLs are included (Fig. 4); the same is true for the correlation between observed and predicted genetic correlations (*r_Q_*) (Fig. 5a,b). Second, bias in the predicted genetic correlations is less when all QTLs are used than when just GWS QTLs are used. Finally, the proportion of CWS QTLs that have effects in the direction of the species difference is similar to that for GWS QTLs and much greater than 0.5 (Table S8).

## ACKNOWLEDGEMENTS

We would like to thank members of the Rausher lab for feedback and advice on the experimental design, analyses, and manuscript; Fred Nijhout for use of the microscope for imaging nectaries. We would particularly like to thank Wendy Dong, Melissa Baldino, Avery Fulford, and Cristian Tolento and members of the Duke Greenhouse Staff, who all helped with plant care and maintenance. This research was supported in part by NSF grant DEB 1542387.

## AUTHOR CONTRIBUTION

ITL and MDR designed the project, collected and analyzed the data, and wrote the manuscript. JLR performed final computational analyses for the linkage map and edited the manuscript. GC annotated the draft genome necessary for the randomization test.

## DATA AVAILABILITY

Raw sequence data from this study are found NCBI Sequence Read Archive [accession: PRJNA732507]. Scripts for sequence processing and linkage mapping are openly available on GitHub at https://github.com/joannarifkin/Ipomoea_QTL/. All other scripts are openly available on GitHub at https://github.com/itliao/IpomoeaNectarQTL.

## SUPPORTING INFORMATION

**Fig. S1** Genetic map and distribution of the 6056 markers used for the QTL analysis.

**Fig. S2** Cluster diagrams based on covariance-component genetic correlations.

**Fig. S3** Frequency distributions of 13 phenotypic traits

**Fig. S4** Pairwise correlations and regressions for the three measured nectar traits and total sugar content.

**Fig. S5** Comparison of correlations derived from RIL means and from variance-covariance (var-covar) components.

**Fig. S6** Pairwise trait correlations among the 13 traits.

**Fig. S7** QTL plots for individual traits.

**Methods S1** Materials and growing conditions and phenotyping nectar traits (nectar volume, nectar sugar concentration, nectary size).

**Methods S2** Summary protocol for ddRAD sequencing.

**Methods S3** Sequence processing and linkage mapping.

**Methods S4** Calculating genetic correlations and broad-sense heritabilities.

**Methods S5** Permutation test for 677 cluster analysis.

**Methods S6** Randomization test.

**Methods S7** Quantifying QTL overlap and permutation test.

**Methods S8** Predicting genetic correlations from QTL properties and permutation analysis.

**Methods S9** Fraser v-test statistic.

**Results and Methods S1** Ascertainment bias analysis and results.

**Table S1** List of individuals phenotyped but NOT genotyped

**Table S2** Genetic correlations derived from variance-covariance components for the 13 phenotypic traits measured.

**Table S3** QTL characteristics for all trait QTLs.

**Table S4** Summary of aggregate QTL characteristics for individual traits

**Table S5** Pairwise QTL overlap.

**Table S6** Randomization test (random placement of QTLs in genome) of whether QTLs within modules are spatially clustered in the genome.

**Table S7** Average predicted genetic correlations, rQ, within and between modules with standard errors (in parentheses).

**Table S8** Number of QTLs with effects in same direction as or opposite direction from difference between species.

**Table S9** Contra-directional QTLs summary.

## Notes

https://github.com/joannarifkin/Ipomoea_QTL/

https://github.com/itliao/IpomoeaNectarQTL

### Competing Interest Statement

The authors have declared no competing interest.

