## Supporting Information for "Modularity and selection of nectar traits in the evolution of the selfing syndrome in *Ipomoea lacunosa* (Convolvulaceae)"

The following Supporting Information is available for this article:

**Fig. S1** Genetic map and distribution of the 6056 markers used for the QTL analysis.

**Fig. S2** Cluster diagrams based on covariance-component genetic correlations.

**Fig. S3** Frequency distributions of 13 phenotypic traits.

**Fig. S4** Pairwise correlations and regressions for the three measured nectar traits and total sugar content.

**Fig. S5** Comparison of correlations derived from RIL means and from variance-covariance (var-covar) components

**Fig. S6** Pairwise trait correlations among the 13 traits.

**Fig. S7** QTL plots for individual traits.

**Methods S1** Materials and growing conditions and phenotyping nectar traits (nectar volume, nectar sugar concentration, nectary size).

**Methods S2** Summary protocol for ddRAD sequencing.

**Methods S3** Sequence processing and linkage mapping.

**Methods S4** Calculating genetic correlations and broad-sense heritabilities.

**Methods S5** Permutation test for cluster analysis.

**Methods S6** Randomization test.

**Methods S7** Quantifying QTL overlap and permutation test.

**Methods S8** Predicting genetic correlations from QTL properties and permutation analysis.

**Methods S9** Fraser *v*-test statistic.

**Results and Methods S1** Ascertainment bias analysis and results.

**Table S1** List of individuals phenotyped but NOT genotyped

**Table S2** Genetic correlations derived from variance-covariance components for the 13 phenotypic traits measured.

**Table S3** QTL characteristics for all trait QTLs.

**Table S4** Summary of aggregate QTL characteristics for individual traits

**Table S5** Pairwise QTL overlap.

**Table S6** Average predicted genetic correlations, *r_Q_*, within and between modules with standard errors (in parentheses).

**Table S7** Randomization test (random placement of QTLs in genome) of whether QTLs within modules are spatially clustered in the genome.

**Table S8** Number of QTLs with effects in same direction as or opposite direction from difference between species.

**Table S9** Contra-directional QTLs summary.

.

**Fig. S1** Genetic map and distribution of the 6056 markers used for the QTL analysis.

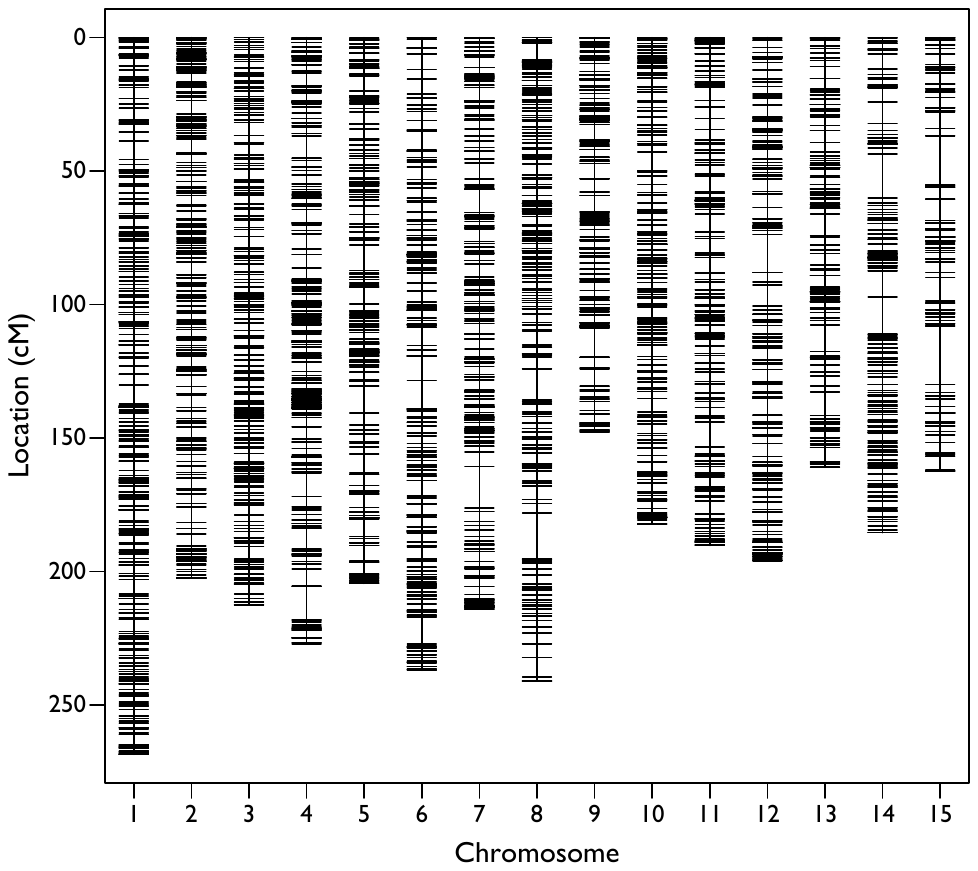

**Fig. S2** Cluster diagrams based on covariance-component genetic correlations. Boxes indicate the three clusters based on hierarchical clustering methods: a) Complete and b) Ward. The three clusters (left to right) – seed (orange), nectar (maroon red), flower size (purple) – are consistent with those shown in Figure 2.

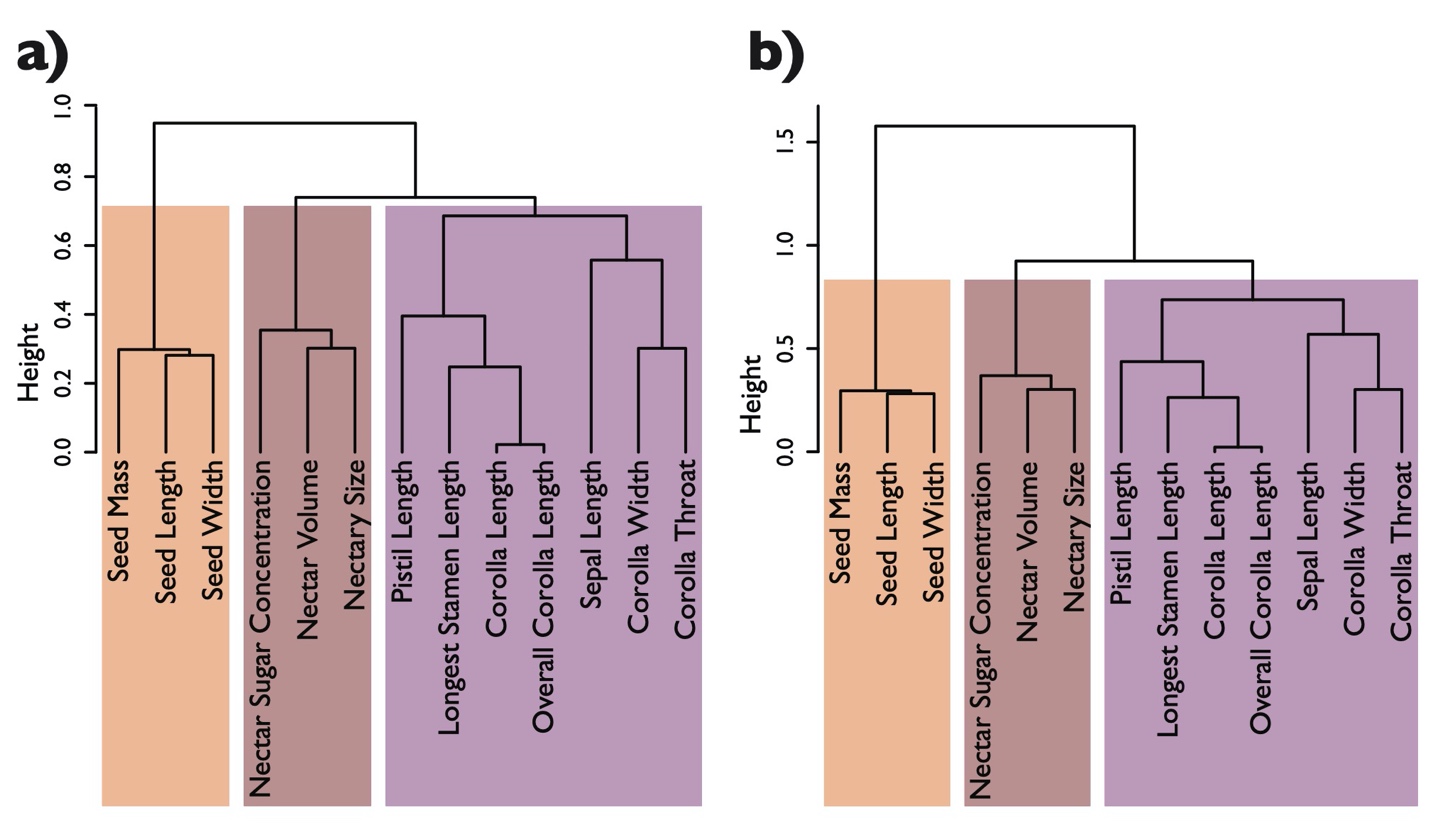

**Fig. S3** Frequency distributions of 13 phenotypic traits. Within each panel, distributions are for *I. lacunosa* selfed offspring (gray), *I. cordatotriloba* selfed offspring (pink), and F5 RILs (purple). Dotted lines indicate the averages within each group, which are shown in Table 1.

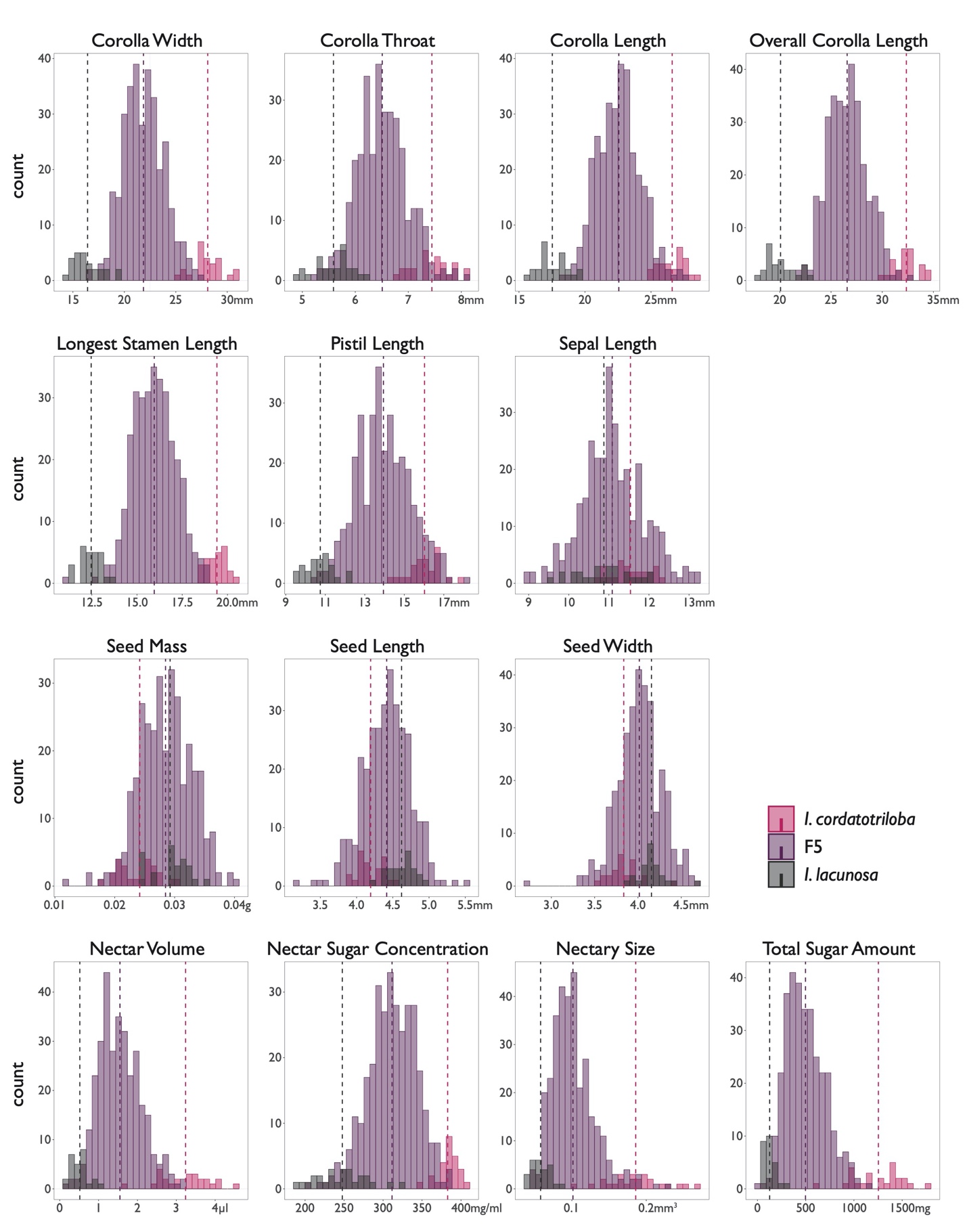

**Fig. S4** Pairwise correlations and regressions for the three measured nectar traits and total sugar content. a) Phenotypic correlations for all 635 F5 individuals. b) Genetic correlations for the 313 RIL means. c) Heatmap of covariance-component genetic correlations based on hierarchical clustering method “mcquitty.” d) Quadratic regression between nectary size and nectar volume. e) Quadratic regression between nectary size and total sugar amount. In d) and e), linear and quadratic coefficients are significant at P < 0.05. f) Linear regression between nectary size and nectar sugar concentration. Blue line: best fit linear regression. Red line: predicted relationship obtained by dividing the regression equation for nectar volume (panel d) by the regression equation for total sugar content (panel e).

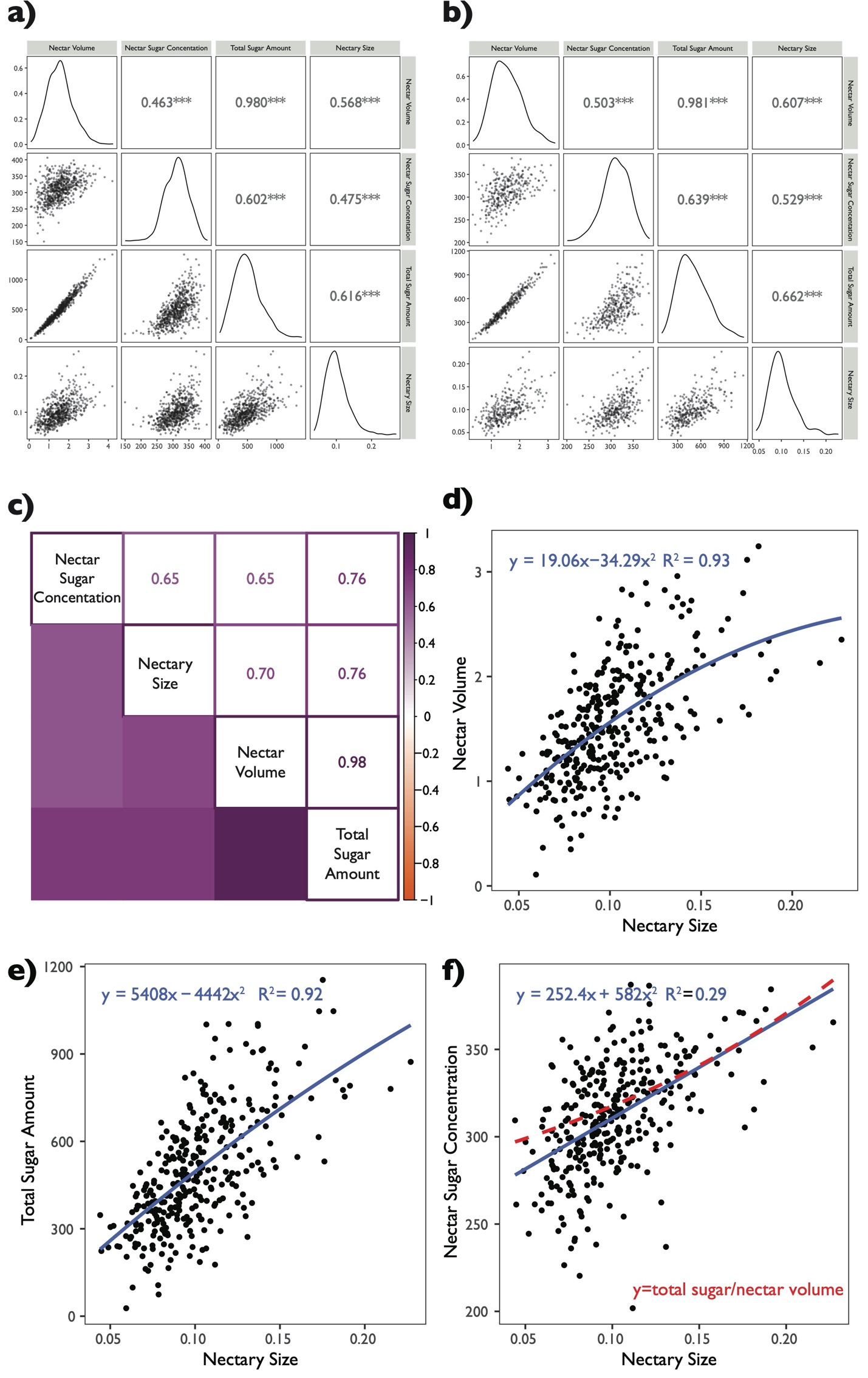

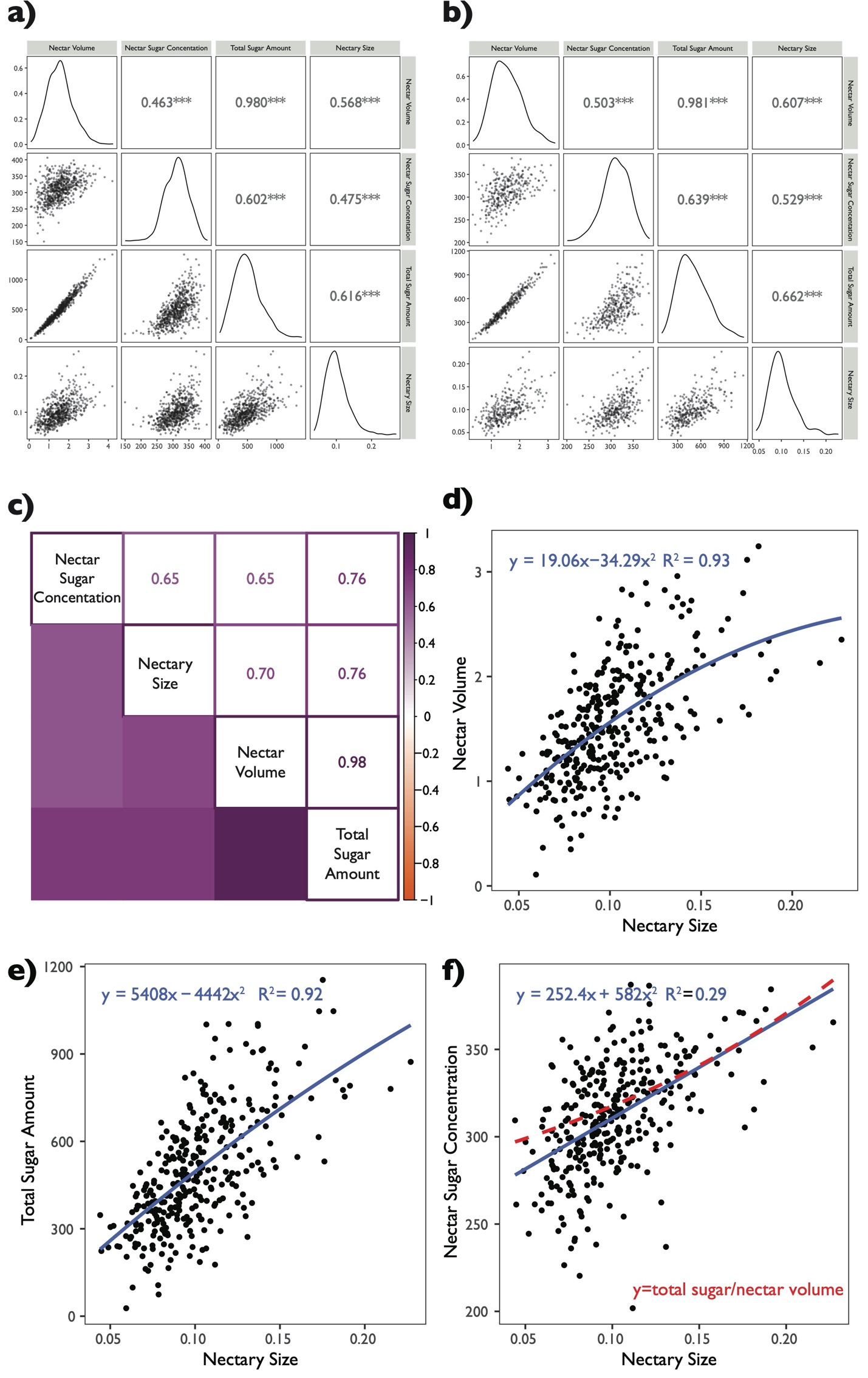

**Fig. S5** Comparison of correlations derived from RIL means and from variance-covariance (var-covar) components. Variance-covariance components are derived from a MANOVA analyses with “Line” as the main effect. Pearson r = 0.981, P < 0.05.

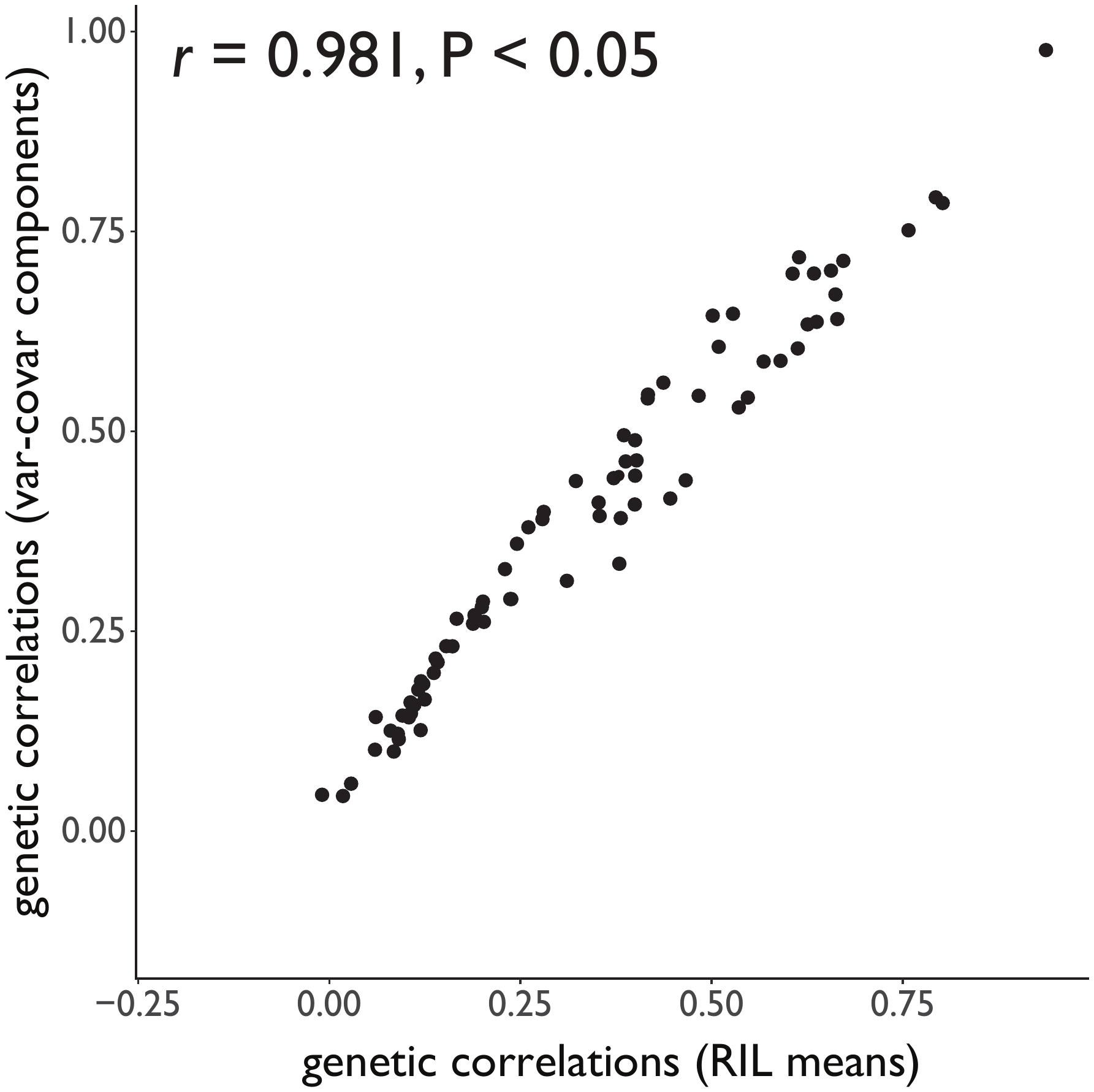

**Fig. S6** Pairwise trait correlations. Lower left triangle displays the scatterplot of points between traits; upper right triangle displays the correlation coefficients and its significance from zero. Panels on the main diagonal show frequency distributions of each trait. a) Phenotypic correlations of all 635 F5 individuals measured for the study. b) Genetic correlations of the 313 RIL means.

a) Phenotypic correlations

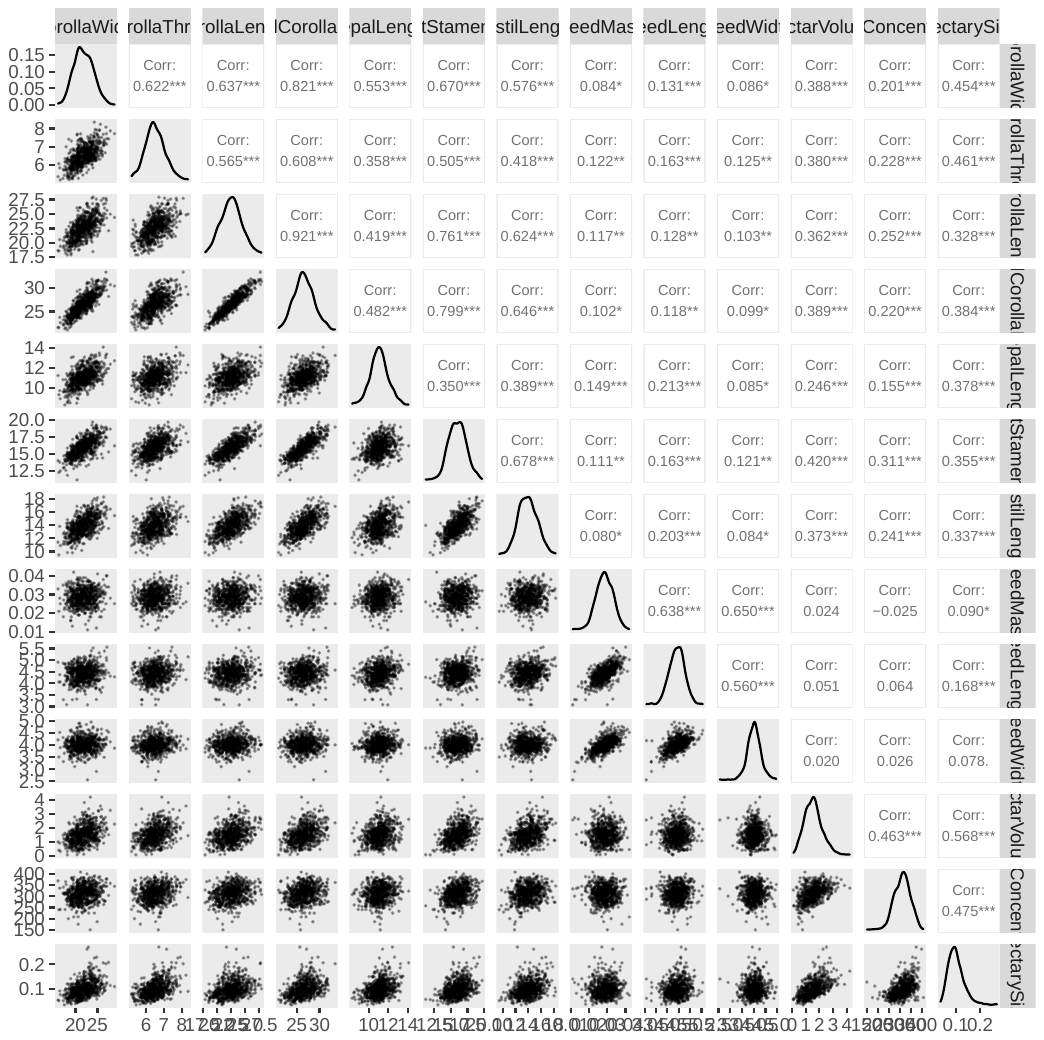

b) Genetic correlations

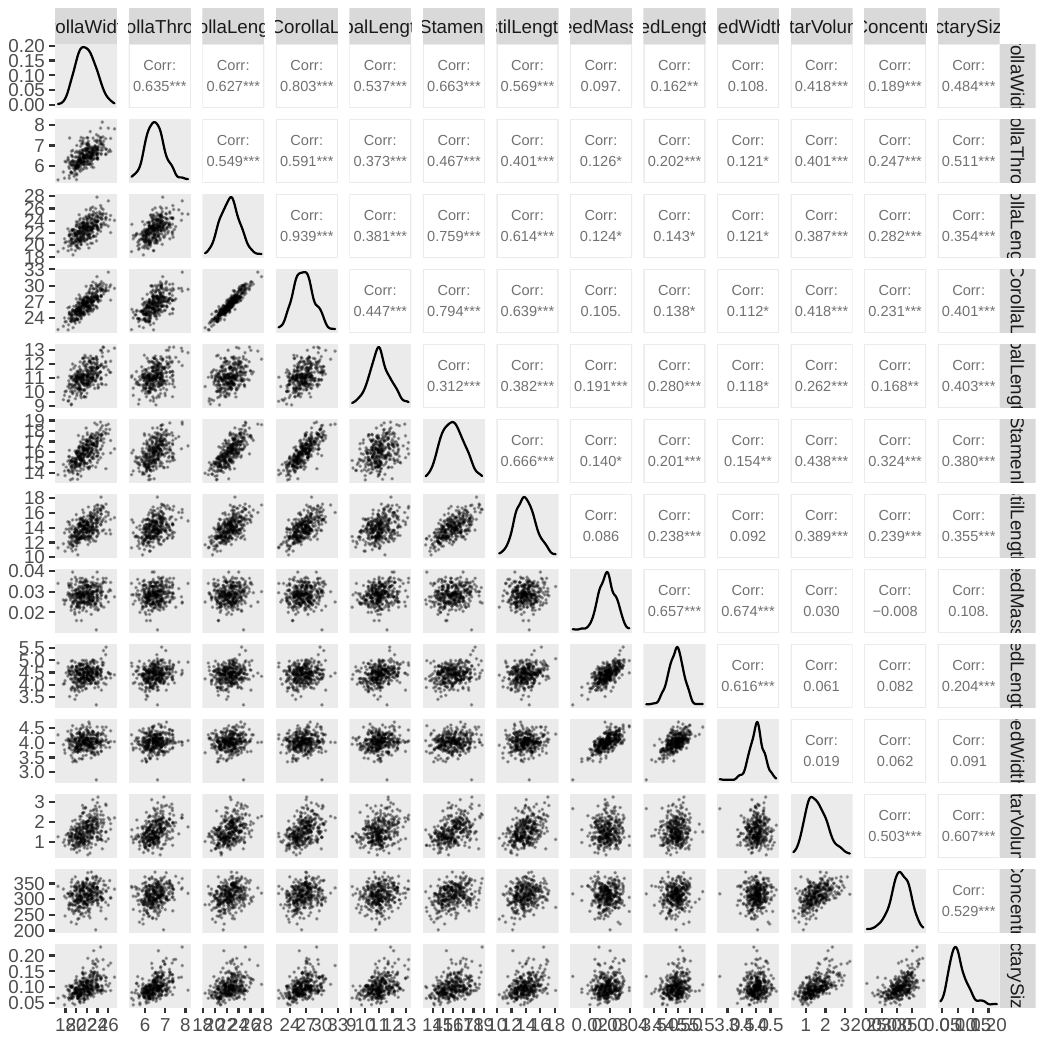

**Fig. S7** QTL plots for individual traits. Solid horizontal lines indicate genome-wide significance threshold determined from 1000 permutations at a significance level of ⍺ = 0.05; peaks above the threshold are determined to be significant. Dashed horizontal lines indicate average chromosome-level significance thresholds determined from 10,000 permutations at a significance level of ⍺ = 0.05. Panels on the left are from analysis with *R/qtl*: dotted line represents a genome scan from the scanone, Haley-Knott regression model; solid black line (with a label indicating the chromosome and position of the peak) represents the result from the multiple QTL, stepwiseqtl model. Panels on the right are from analysis with *R/qtl2* LOCO model.

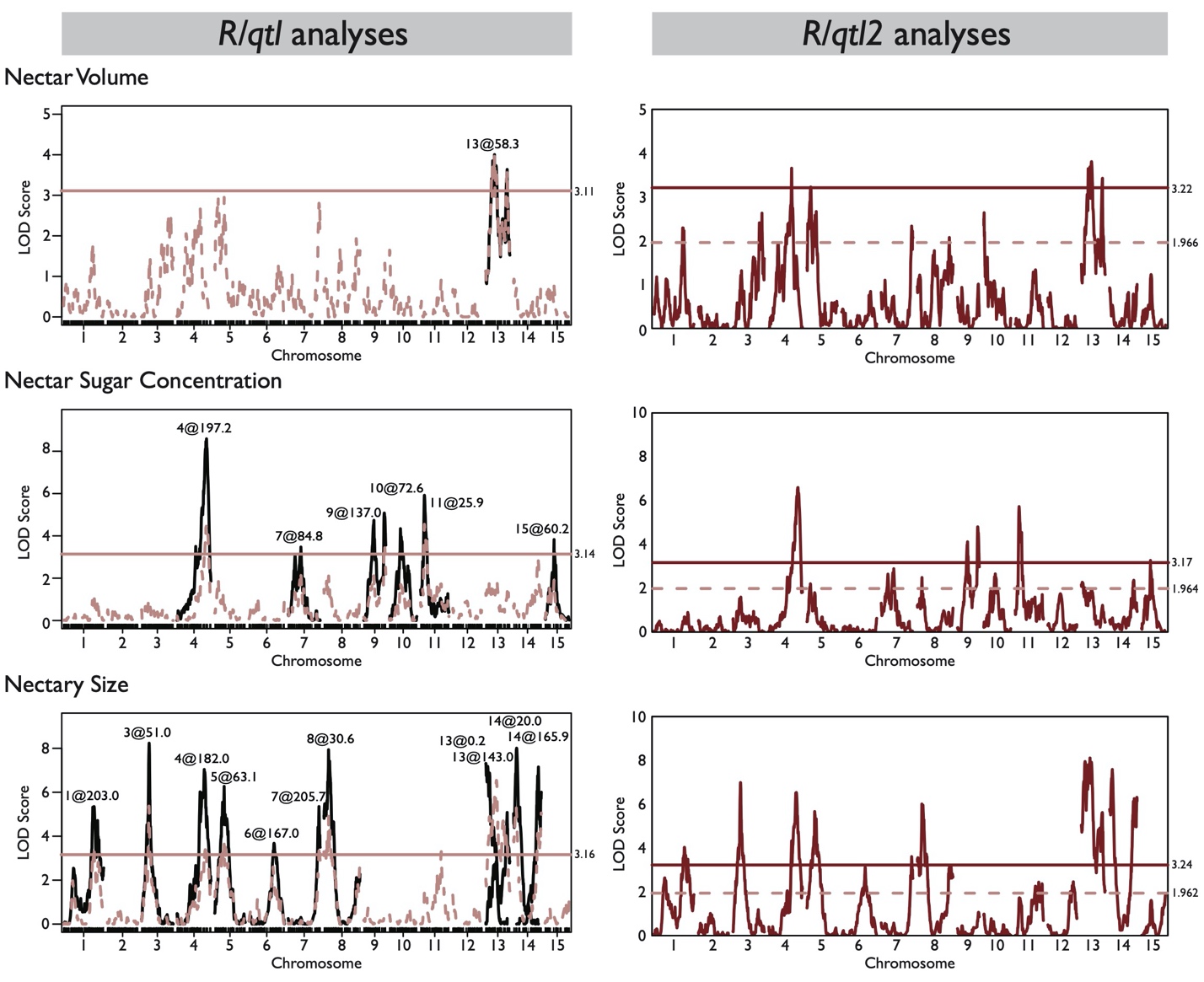

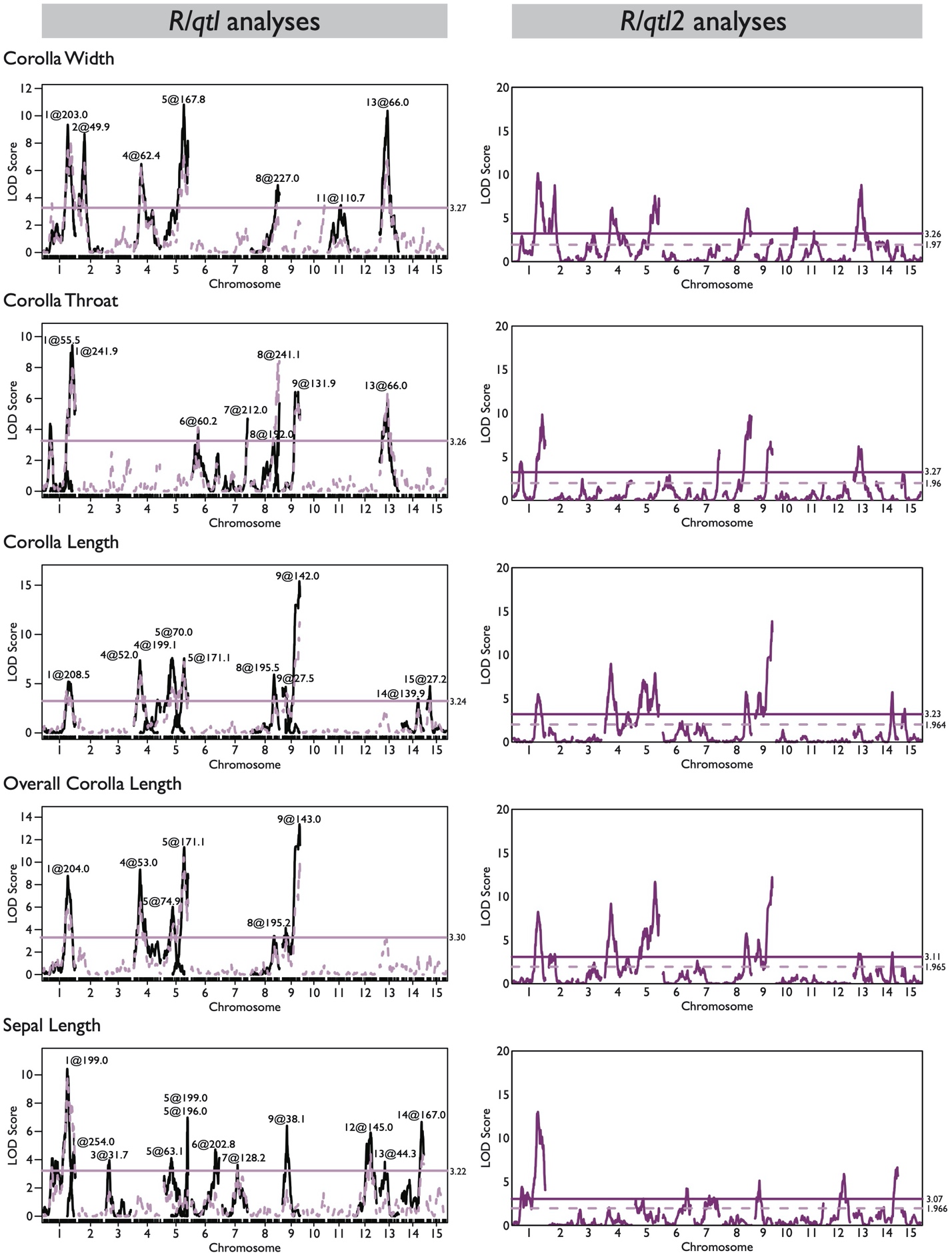

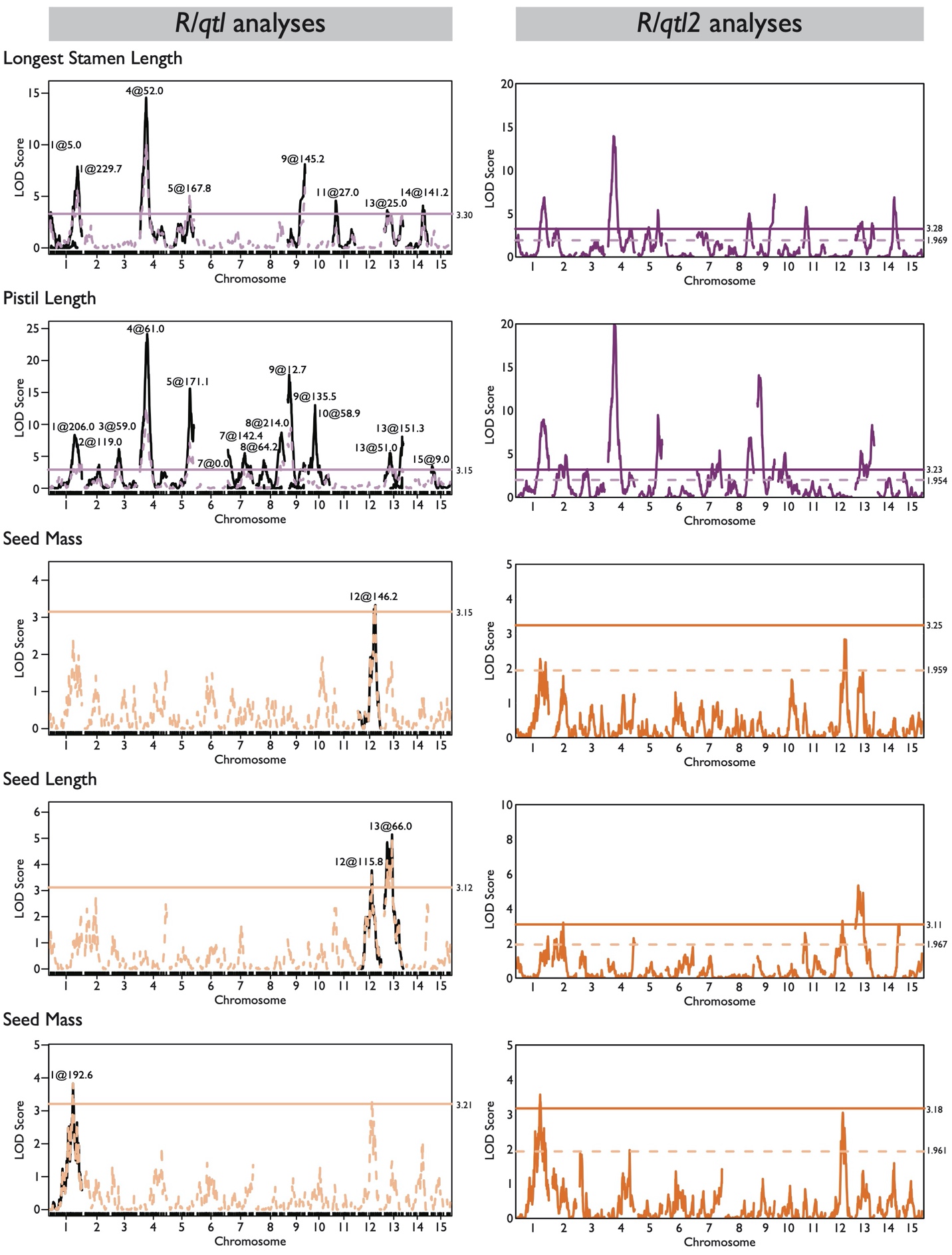

**Methods S1** Materials and growing conditions and phenotyping nectar traits (nectar volume, nectar sugar concentration, nectary size).

*Materials and growing conditions*

One *I. cordatotriloba* individual from Conway, South Carolina (33.94713, -79.01940; hereafter C parent) was crossed by one *I. lacunosa* from Kinston, North Carolina (35.23971, -77.57392; hereafter L parent) to generate F1 hybrids (Duncan & Rausher, 2013; Rifkin *et al.*, 2021). One F1 (CL5) was selfed to generate a mapping population of 500 F2s; these were selfed by single seed decent for 3 more generations to generate 322 recombinant inbred lines (RILs) at the F5 generation. About 37% of the original F2s planted (186/508) did not make it to the F5 generation due to poor germination, flowering, and/or inviable seed set.

Four individual seeds per RIL and 50 individual selfed seeds from the parent lines (L parent, C parent) were planted for a total of approximately 1400 seeds. Approximately 2 individuals per line and 24-25 individuals per each parent line were selected for genotyping and phenotyping.

Seeds were scarified by cutting off a small part of the seed coat to induce quick germination. Scarified seeds were planted directly into soil (Farfard 4P) in 24-cell seed packs in a randomized fashion. Plants were grown under 12-hour light:12-hour dark in 21-23ºC­ to induce floral bud formation. Trays of seedlings were rotated around the growth chamber to minimize micro-environmental effects. After a month, the plants were transferred to the Duke University Greenhouses, where the plants were grown under the following conditions: 12-hour light:12-hour dark, light temperatures 23-26ºC, dark temperatures 16-19ºC, 65% relative humidity & 700umol/s/s/cm2 light intensity. Plants were watered daily and fertilized with 300ppm N of 10-30-20 once a week.

*Nectar trait phenotyping*

As a rule, the first flower produced by a plant was not measured for any trait. For most individuals (675 out of 686 total individuals), floral and nectar traits were measured on at least 3 flowers per individual plant. All measurements were performed between 9 AM to 12 PM from October 2017 – May 2018. Floral size traits and nectar traits were measured on alternating days because flowers needed to be capped in order to measure nectar accurately, and this may have distorted final flower form.

Methods for measuring nectar volume and sugar concentration are detailed in Rifkin *et al.*, 2019a and summarized here: the day before the flower opened, the flower bud was capped with a straw covered with parafilm on one end to limit evaporation. The following day, nectar volume was collected using 2µl microcapillary tubes (Drummond Scientific), and the height of the liquid in the tubes was measured with a caliper. All nectar was expelled onto a Master‐53M ATAGO refractometer to measure sugar concentration. Because nectar sugar concentration readings are often imprecise for low amounts of liquid, 2.5µl water was added, and a second sugar concentration reading was taken. Nectar volume was converted into µl using this equation: V = 2 μl*(height of nectar in tube/32 mm). Nectar sugar concentration readings were converted as suggested by Bolten *et al.*, 1979. First, Brix readings (weight/weight percentages; w/w) were converted into mg/ml values using this equation: (mg/ml = 0.0524(w/w)^2^ + 9.6554(w/w) + 1.3904). For the dilutions, values were adjusted using the equation: (actual nectar amount + 2.5 μl)/actual nectar amount. The average of the two values was used for further analyses. Total sugar amount was calculated as the product of nectar volume and nectar sugar concentration.

Nectaries from the individuals that were measured for nectar volume and sugar concentration were collected for measurement and stored in 30 µl RNALater in a -20ºC freezer before being imaged. Stored nectaries were photographed using a Leica MZFLIII stereomicroscope set at 1X magnification and connected to a 4D HD digital microscope camera from Microscope Systems, Inc. (https://microscope-systems.com/). The images were calibrated with a microscope calibration ruler slide and found to be 345 pixels per 1 cm. Nectaries were measured from images using the imaging software on FIJI (Schindelin *et al.*, 2012) by first selecting 6 points to designate the outline of the nectary (Fig. 1c). The distance between certain sets of points was calculated to obtain the variables required to estimate the nectary size, which is modeled after the volume of a frustrum using the equation:

$$V= \frac{\pi h}{3}\left[ \left( r_{1}^{2}+r_{1}r_{2}+r_{2}^{2} \right)-\left( r_{3}^{2}+r_{3}r_{4}+r_{4}^{2} \right) \right]$$

where *h* is the nectary height, *r_1_* is the nectary top outer radius, *r_2_* is the nectary top inner radius, *r_3_* is the nectary bottom outer radius, and *r_4_* is the nectary bottom inner radius (Fig. 1d). Because the bottom inner radius was difficult to measure from the image alone, it was assumed that the nectary thickness (taken as the difference between the top outer and top inner radiuses) tapered off from the top to the base of the nectary and was assumed to be non-existent at the base of the nectary. Thus, the bottom inner radius was the same as the bottom outer radius. Raw images, raw and cleaned data files, and scripts are found on Dryad.

**Methods S2** Summary protocol for ddRAD sequencing.

We used a modified version of the Peterson double-digest restriction assisted DNA (ddRAD) library preparation protocol (Peterson *et al.*, 2012) combined with the Rieseberg lab genotype-by-sequencing protocol (Ostevik, 2016; a detailed description of the protocol can be found here: https://github.com/itliao/IpomoeaNectarQTL/blob/main/ddRAD_protocol/ddRAD_Protocol_180916.docx). Briefly, samples were digested with MspI and EcoRI. EcoRI and MspI adapters with specific 8-base barcodes were ligated to the samples. Each plate (largest number of samples – 96; a total of 8 plates of samples) was pooled and cleaned using AMPure beads (Beckman Coulter). Cleaned, pooled samples were quantified before undergoing PCR amplification with Illumina-specific barcode adaptors for each plate. PCR products were cleaned with AMPure beads, quantified, and all samples pooled in equal amounts (100 ng) together before sequencing over 4 lanes of Illumina HiSeq 4000 (150bp PE) reads at the Duke Center for Genomic and Computational Biology Sequencing and Genomic Technologies Core. Of the 322 lines, two individuals were genotyped for 294 RILs, but only one for 28 RILs.

**Methods S3** Sequence processing and linkage mapping.

Detailed methods can be found in Rifkin *et al.*, 2021; scripts can be found on Github: https://github.com/joannarifkin/Ipomoea_QTL/.

Briefly, raw sequence reads were demultiplexed using sabre (<https://github.com/najoshi/sabre>), and then aligned to the *I. lacunosa* draft genome (approximately 50 scaffolds) using NextGenMap v0.5.5 (Sedlazeck *et al.*, 2013). Aligned reads were then sorted and cleaned using PicardTools. We used Lep-Map3 (Rastas, 2017) to call genotype probabilities, relate the genotypes back to the parental genotypes, and assigned to 15 linkage groups. F5 RIL markers were ordered with Lep-Map3 and related back to the draft genome assembly with Lep-Anchor (Rastas, 2020).

**Methods S4** Calculating genetic correlations and broad-sense heritabilities.

Genetic and environmental correlations were calculated for all traits on F5 RILs in two ways. In one method, the genetic correlations were estimated as the correlations of line means (Falconer & Mackay, 1996). The second method estimates genetic correlations from variance and covariance components in a Multivariate Analysis of Variance, in which line is the main effect (Falconer & Mackay, 1996). These components were calculated using the *car* package in R (Fox & Weisberg, 2019). To obtain the genetic correlation for a pair of traits, the between-line Sum of Products was divided by the associated degrees of freedom to obtain the between-line Mean Cross Product (MCP). The error (within line) MCP was subtracted from the between-line MCP to obtain a value that is *n* times the between-line covariance component, *Cov_G_*, where *n* is a function of the number of replicate individuals per line. Similarly, between-line variance components were obtained by subtracting the within-line mean square from the between-line mean square, which again produces a value that is *n* times the between-line variance component, *Var_G_*. The genetic correlation was then calculated for traits x and y as

$r_{G}= \frac{n {Cov}_{G}}{\sqrt{n{Var}_{G}\left( x \right) n {Var}_{G}\left( y \right)}}= \frac{{Cov}_{G}}{\sqrt{{Var}_{G}\left( x \right) {Var}_{G}\left( y \right)}}$

(Falconer & Mackay, 1996). Because these correlations are calculated from inbred lines, they represent broad-sense genetic correlations. Correlation matrices were plotted using *corrplot* package (Wei & Simko, 2017).

Broad-sense heritabilities were calculated for each trait in a similar fashion as

$H^{2}= \frac{{Var}_{G}}{{Var}_{G} + {Var}_{E}}$ ,

where *Var_E_* is the within-line variance component, equal to the error mean square. We determined the variance components of each trait using the *lme4* package (Bates *et al.*, 2015) and the *lmer* function by treating “Line” as a random effect. The significance was calculated using the *lm* function and performing an ANOVA test using the *car* package (Fox & Weisberg, 2019).

Because the two methods of calculating genetic correlations give very similar results (see Text), we use the variance-covariance component correlations for all analyses.

To understand whether nectary size is proportional nectar volume and nectar sugar amount, we first calculated the total sugar amount by multiplying the measured values of nectar volume and nectar sugar concentration. We performed quadratic regressions between nectary size and nectar volume and between nectary size and total sugar amount. We divided these two regressions (total sugar amount by nectary size/nectar volume by nectary size) to derive the expected regression the between nectary size and sugar concentration. We then compared this derived value with the actual regression between nectary size and the measured value for nectar sugar concentration.

**Methods S5** Permutation test for cluster analysis.

As an indication of whether the identified modules are real, we performed a permutation test. For each permutation, we randomly assigned each observed genetic correlation to a random pair of traits. We then calculated the average of all correlations that represent within-module trait pairs and the average of all correlations that represent between-module trait pairs and computed the difference between the two averages. We ran 1,000 permutations, yielding 1,000 values of this difference. Finally, we compared the distribution of these values to the observed difference. The proportion of differences from the permutation trials that are greater than or equal to the observed difference indicates the probability that the observed level module differentiation would occur by chance.

**Methods S6** Randomization test.

To determine whether the overall degree of QTL overlap within modules was greater than expected by chance, we performed a randomization test, in which we assigned the actual QTLs to random positions in the genome, calculated QTL overlap for all pairs of traits, then averaged the overlap for pairs of traits that occur in the same module. When placing the QTLs in random positions, we maintained the size of the QTL and took into account gene density, reasoning that a QTL is more likely to occur in regions of high gene density than regions of low density. Using our annotated genome of *I. lacunosa*, we determined the number of genes occurring in 50kb bins across the genome. To assign a position to a QTL, we first chose a chromosome randomly in proportion to the total number of genes on that chromosome. We then chose a 50kb bin on that chromosome randomly in proportion to the number of genes in different bins. Finally, within the chosen bin, we selected a position randomly. We ran 1,000 permutations and calculated the average QTL overlap within modules for each permutation. The probability that the observed average QTL overlap within a module can be explained by random placement is the proportion of permutations that have a higher average QTL overlap than that observed. We ran this test for each module separately and an overall test on all within-module trait pairs.

**Methods S7** Quantifying QTL overlap and permutation test.

We quantified the degree of QTL overlap between two traits using the Jaccard index:

$$QTL overlap=\frac{n_{12}}{n_{1}+n_{2}-n_{12}}$$

where *n*_1_ is the number of trait 1 QTLs, *n*_2_ is the number of trait 2 QTLs, and *n*_12_ is the number of QTLs that have overlapping QTL confidence intervals for trait 1 and trait 2. To determine how well QTL overlap within and between modules explains the observed modularity, we calculated the mean and standard errors of QTL overlap within and between modules. We tested whether there was greater overlap within modules than between modules using a permutation test in which QTLs were assigned randomly to modules.

To assess how well QTL overlap explains the observed genetic correlations, we calculated the correlation between genetic correlation values and QTL overlap values. We assessed whether the correlation was significant by a permutation test in which we permuted the values of the genetic correlations without changing the QTL overlap values.

**Methods S8** Predicting genetic correlations from QTL properties and permutation analysis.

We also assessed the degree to which QTL effects, particularly those from co-localizing QTLs, can predict genetic correlations between traits, using a modification of the approach in Gardner and Latta (2007). We treated the F5 individuals as inbred lines, in which each line is homozygous for one of the alleles at a particular QTL. For each QTL for a particular trait *i*, we assigned lines homozygous for the *I. cordatotriloba* allele a value of ½ the relative homozygous effect (RHE), symbolized by *E_i_*, and lines homozygous for the *I. lacunosa* allele a value of -½ RHE, symbolized by –*E_i_*. For QTLs that overlap for two traits *i* and *j*, we calculated the genetic covariance of individual QTLs as

${Cov}_{{QTL}_{ij}}=p_{1}\left( E_{i}- \bar{E_{i}} \right)\left( E_{j}- \bar{E_{j}} \right)+(1-p_{1})\left( {-E}_{i}- \bar{E_{i}} \right)\left( -E_{j}- \bar{E_{j}} \right)$

Where *p*_1_ is the proportion of lines homozygous for the *I. cordatotriloba* allele, and

$\bar{E_{i}}=p_{1} E_{i}+ (1-p_{1})(-E_{i})$.

Similarly, the genetic variance of each individual QTL for a trait *i* (including those that do and do not overlap another QTL) were calculated as

${Var}_{{QTL}_{i}}= p_{1}\left( E_{i}- \bar{E_{i}} \right)^{2}+(1-p_{1})\left( {-E}_{i}- \bar{E_{i}} \right)^{2}$

The predicted genetic covariance between two traits *i* and *j* is then just the sum of the covariances of the individual QTLs:

${Cov}_{G_{ij}}= \sum_{QTL} {Cov}_{{QTL}_{ij}}$

and the predicted genetic variance of a trait *i* is the sum of the variances of the individual QTL variances for that trait:

${Var}_{G_{i}}= \sum_{QTL} {Var}_{{QTL}_{i}}$

Finally, the predicted genetic correlation between two traits is

$r_{Q_{ij}}= \frac{{Cov}_{G_{ij}}}{\sqrt{{Var}_{G_{i}} {Var}_{G_{j}}}}$

We designate this correlation with a subscript *Q* to indicate it is calculated from QTL properties and is to be distinguished from the estimated genetic correlations $r_{G_{ij}}$ calculated from variance and covariance components.

These calculations assume that the locations of overlapping QTLs are identical, which means that the QTLs for the two traits have the same value of *p_i_*. For a particular QTL, we used the value of *p_i_* corresponding to marker with the highest LOD score. In practice, this means that overlapping QTLs will not have exactly the same value of *p_i_* because the peaks will not necessarily correspond to the same marker. Therefore, as an approximation to the joint *p_i_* of two traits, we identified the midpoint of the overlap interval and used the allele frequencies from the marker closest to the midpoint.

We assessed how well genetic correlations predicted from QTL properties matched the estimated true genetic correlations using correlation analysis. Specifically, we calculated the Pearson correlation coefficient for the correlation between the true genetic correlations and the predicted genetic correlations. Because the predicted genetic correlations may inherently reflect the QTL overlap (covariances could only be calculated between QTLs that overlap), we calculated the correlation between the predicted genetic correlations and QTL overlap. We also calculated analogous correlations between genetic correlations and bias, which is the difference between the predicted and true genetic correlations, *r_Q_*−*r_G_* (Gardner & Latta, 2007). Finally, as an indicator of the “completeness” of how our QTL identification affects predicted correlations, we calculated the correlation between bias and average total RHE for a trait pair, where total RHE is the sum of the RHE values of individual QTLs for a trait.

To determine the significance of these correlations, we performed a permutation analysis. For each, permutation, we reassigned QTLs to random traits, calculated *r_Q_*, bias and average total RHE, and then calculated the Pearson correlation coefficients described above. The proportion of 1,000 permutations with correlations greater than the observed correlation is taken as the probability of obtaining the observed correlation by chance. We also considered two scenarios: (1) pairwise correlations among all 13 traits, and (2) pairwise correlations of only floral and nectar traits. Finally, we determined whether bias involving floral and nectar traits differed for the two QTL sets by bootstrapping (1,000 replicates).

**Methods S9** Fraser *v*-test statistic.

The *v* test, which we also refer to as the Fraser test, calculates the tests statistic

$v= \frac{\sigma_{b}^{2} - \frac{\sigma_{p1}^{2}}{n_{p1}} - \frac{\sigma_{p2}^{2}}{n_{p2}}}{\sigma_{R}^{2} H^{2} c}$	,

where $\sigma_{b}^{2}$is the between-species variance component among the cross parents, $\sigma_{pi}^{2}$ is the phenotypic variance among parent species *i*, $n_{pi}$ is the number of parent species *i*, $\sigma_{R}^{2}$ is the phenotypic variance among the RILs, *H*^2^ is the broad sense heritability, and *c* is a constant equal to 1.0 for RILs (Fraser, 2020). Variances and covariances were calculated using appropriate analyses of variance as implemented by the Varcomp procedure in the SAS statistical software package version 9.4 (SAS Institute, 2013). Significance of *v* is based on the cumulative F distribution with (1, *k ─* 1) degrees of freedom, where *k* is the number of RILs (Fraser, 2020). We used a sequential Bonferroni correction (Holm, 1979) to correct significance levels for multiple comparisons.

**Results and Methods S1** Ascertainment bias analysis and results.

We examined the possible effects of ascertainment bias on our inferences of selection based on the QTL-EE and Fraser tests. Anderson and Slatkin (2003) demonstrated that ascertainment bias can greatly increase the probability of false positives in a QTL-EE sign test. However, the ascertainment bias they modeled was as extreme as possible: they assumed that researchers chose the one trait showing the greatest between-species difference out of *N* candidate traits. While this type of choice might be appropriate for examining molecular and physiological characters (e.g. Fraser, 2020), we believe it does not reflect how researchers studying divergence in organismal traits choose characters to study. Usually, traits are chosen because they are of evolutionary interest, not because they are the most diverged trait out of a predefined set of traits. For example, someone interested in mating system divergence would naturally choose floral traits, even though the species in question may have diverged more in, say, root characters than in floral characters. In choosing traits to study in this manner, there is no reason to believe that researchers are choosing those that have diverged to the greatest degree. These considerations thus raise the question of the extent to which ascertainment bias of this type increases the probability of false positives in the sign test.

To address this question, we have conducted a set of simulations analogous to those of Anderson and Slatkin, but with different assumptions about how traits are chosen. We examine three different criteria for choosing a trait: (1) choose the most divergent trait from *N* candidate traits (the Anderson and Slatkin method); (2) choose randomly from a set of *N* candidate traits; and (3) choose randomly from the 50 percent of *N* candidate traits with the highest divergence. Criterion (2) and (3) are meant to be more representative of how evolutionary biologists studying organismal divergence choose traits to study. Criterion (2) is the least conservative in assuming that traits can be chosen for study even if they show very little divergence relative to other traits. Criterion (3) is more conservative in assuming that traits showing the least divergence are passed over; however, if divergence is above a certain threshold, a trait can be chosen regardless of its degree of divergence compared to other traits above that threshold. We designate the threshold as β, which in this case was 0.5. More generally, for a given threshold β, traits are ordered from highest to lowest divergence, and the lowest (1 – β)*N* traits are discarded. The study trait is then drawn randomly from the remaining traits.

Simulations were performed for traits controlled by either 10 or 11 loci, each with an equal effect on the character. For a given trial with 10 loci, *N* “traits” were simulated by randomly assigning alleles 1 or 2 to each of 10 loci for each trait, where *N* ϵ {1, 10, 25, 50, 100}. “1” alleles are assumed to have effects in the opposite direction of the species difference, while “2” alleles are assumed to have effects in the same direction as the species difference. Each allele was envisioned as replacing a “0” allele at each locus, and thus causing a divergence in the trait. For cases in which there were more “1” alleles than “2” alleles, the alleles were switched (i.e. “1” was changed to “2” and “2” was changed to “1”) to preserve the proper direction of trait change.

For each trait, the number of “2” alleles was counted, and the *N* traits were ordered from highest to lowest number of “2” alleles. This ordering corresponds to ordering the traits in terms of divergence. Three methods of choosing the study trait were then implemented: (1) choose the trait with the highest number of “2” alleles (“Top 1” method), which is equivalent to choosing the most diverged trait and corresponds to the method of Anderson and Slatkin; (2) choose randomly among the traits (“Random” method), which corresponds to choosing randomly among all diverged traits; and (3) choose randomly among the top 50% (corresponding to β = 0.5) of the traits (“Random50” method), which implements a bias against choosing traits that diverged less than the average amount.

For each *N*, this process was repeated 10,000 times, and for each replicate the number of “2” alleles was recorded. With no ascertainment bias, under the null hypothesis that divergence at the chosen locus is neutral, obtaining either 9 or 10 “2” alleles at the locus has a probability of 0.0215, and would thus be significant at the P = 0.05 level. Another way of saying this is that the false positive rate (FPR) would be 0.0215. Accordingly, for each of the three methods of choosing we can estimate the FPR as the probability of obtaining either 9 or 10 “2” alleles in the simulations. The ratio of this FPR to the theoretical FPR of 0.0215 indicates the degree to which ascertainment bias inflates the probability of a false positive.

One complication arises with simulations using 10 loci: one possible outcome is 5 “1” alleles and 5 “2” alleles. In such an individual, the positive and negative effects would cancel, and there would be no divergence of the trait. A researcher would thus not choose that trait to study. We adjusted for this complication in the following way: when each of *N* traits were assigned their alleles, we discarded any trait that had 5 “1” alleles and 5 “2” alleles. This procedure produced *N* traits, each of which exhibited some degree of divergence. However, we still considered a trait with 9 or 10 “2” alleles to represent an unlikely occurrence due to chance. In fact, the expected FPR with this adjustment to allele sampling is [1/(1 – prob(5 “1” and 5 “2” alleles)] 0.0215 = 0.0285, which is still less than P = 0.05. Simulations using 11 loci were performed in exactly the same way, except that this adjustment was not performed because all combinations of alleles exhibit some degree of trait divergence. With 11 alleles, the criterion for a false positive with no ascertainment bias and either 10 or 11 “2” alleles, the probability is 0.0117, which is significant at P < 0.05.

As Anderson and Slatkin (2003) found, with the Top 1 method, the FPR increased dramatically as *N* increases (Table A1). In fact, for 10 loci, the FPRs we obtained are very close to those obtained by Anderson and Slatkin and differ only at the 3^rd^ decimal place, as would be expected from stochastic simulations. With 10 loci and *N* = 1, we found estimated FPR to be very close to the theoretical values (0.0290 from simulations vs. 0.0285 theoretical). However, with *N* = 50, the expected FPR increased to 0.765, and with *N =* 100, obtaining a false positive is almost a certainty (FPR = 0.948). These figures represent increases over the FPR for *N* = 1 by factors of 26 and 32, respectively. Similarly, with 11 loci and *N* =1, the expected FPR is 0.0120 (theoretical value = 0.0117). But the FPR increase to 0.451 with *N* =50 and to 0.696 with *N* = 100 (increases by factors of 38 and 59).

By contrast, choosing traits randomly among *N* candidate traits has little effect on the FPR for either 10 or 11 loci (Table A1). This result should not be surprising because choosing randomly among *N* randomly simulated traits should not bias the FPR.

Finally, choosing randomly among the 50 percent of traits showing the most divergence does increase the FPR. However, it does so only by a factor of about 2.0, regardless of *N* and regardless of the number of loci (Table A1, Ratio column). This result indicates that less radical methods for selecting a study trait than Top 1 give much smaller effects of ascertainment bias on FPR. Indeed, if the Random50 method is a reasonable approximation of how traits are chosen for study, one needs to adjust the significance level for the trait only by a factor of 0.5; instead of using P < 0.05 as the criterion for significance, one would use 0.025.

This result suggests that when the study trait is chosen randomly from the β*N* most highly diverged traits (0 < β < 1), the expected FPR should be 1/β times the FPR under completely random sampling (the expected FPR with no ascertainment bias). We term 1/β the “FPR ratio”. We tested whether this was the case using simulations with 11 loci and *N* = 100. We chose the study trait from the fraction β of 100 candidate traits with the highest divergence for β = (0.1, 0.2, 0.3, . . . 0.9, 1.0) and estimated the FPR with 10,000 replicate simulations. There was excellent agreement between the FPR ratio from the simulations and the predicted value (Figure A1).

This relationship indicates a way that one can adjust significance levels to account for ascertainment bias. In particular, if α is the nominal significance level and if it is reasonable to assume that the study trait represents a random choice from the β*N* most diverged traits, then the adjusted significance level is αβ. For example, if α = 0.05 and if it is reasonable to assume that the study trait represents a random choice from the 1/3 most diverged candidate traits, then β = 1/3, and the new adjusted significance level is αβ = 0.05 1/3 = 0.0167. Thus, if the study trait has a nominal significance level that is less than the new adjusted significance level, this would suggest that ascertainment bias was not involved in choosing the study trait. However, if the study trait has a nominal significance level that is greater than the new adjusted significance level, this would suggest that ascertainment bias may have affected the choosing of the study trait.

We applied this analysis to the QTL-EE sign test for GWS QTLs and ALL QTLs. Although there is no theoretical or empirical guidance, we believe it to be an unrealistic assumption that traits chosen for study are among the top 0.15 or less, which implies β > 0.15. Consequently, we used a corrected P value of 0.05 • 0.15 = 0.0075 to determine test significance. Because a β value less than 0.4 may also be unreasonable, we interpret a P value of 0.05 • 0.4 = 0.02 as possibly indicating significance. For GWS QTLs, the test remains significant after correcting for ascertainment bias for corolla width, while corolla length is of borderline significance using 0.0075 as the critical value (Table S4A). With 0.02 as the critical value, corolla throat is of borderline significance, where the β value is 0.627, indicating we cannot rule out the null hypothesis that this trait diverged neutrally. Because none of the other traits are nominally significant by the QTL-EE sign test, correction for ascertainment bias was not performed.

Results for ALL QTLs yield greater significance: for five traits (corolla width, overall corolla length, longest stamen length, pistil length and nectar sugar concentration), the neutral hypothesis can be rejected using the more stringent criterion corresponding to β > 0.15 (Table S4B). Neutrality can probably also be rejected for corolla length (maximum β = 0.19), and possibly for nectar volume (maximum β = 0.352). Ascertainment bias may have played a role in selecting corolla throat (maximum β = 0.44) and nectary size (maximum β = 0.556), and these two traits may have diverged neutrally.

In general, the Fraser test yielded much stronger significant results than the QTL-EE test. Using the criterion of adjusted P value less than 0.0075, all floral and nectar traits except sepal length and pistil length are significant even after a sequential Bonferroni correction (Table A2). We thus believe that ascertainment bias unlikely explains the significance of the Fraser test for these traits.

Another way of assessing the effects of ascertainment bias in the Fraser test was suggested by Fraser (2020). In particular, he suggested that one calculate the maximum number of candidate traits that would be required for a “Top 1” choice to yield the calculated significance level (*N**): 0.05/*p*, where *p* is the significance level. These values are given in Table A2. For all nectar traits and most floral size traits, *N** > 120. It is difficult to imagine that there could be more than 120 nectar and floral candidate traits that we could have chosen among in our study of divergence. We therefore infer that these traits were very likely subject to divergent selection. The cases of corolla throat (*N** = 50) and pistil length (*N** = 25) are less extreme, making a similar inference less credible. It should be noted, however, that this correction is very conservative because it assumes that the examined trait had the largest divergence among the *N** candidate traits.

In summary, these considerations of ascertainment bias suggest that selection contributed to the divergence of most of the flower size and nectar traits examined, with the possible exception of corolla throat, pistil length, and nectary size. By contrast, there is no evidence that selection contributed to divergence in seed characters and sepal length.

Table A1 Probability of false positive under various schemes for choosing study traits. *N* is number of candidate traits. A&S Result is false positive rate found by Anderson and Slatkin (2003) using the Top 1 method. TFPR is the theoretical false positive rate with no ascertainment bias and a significance level of P = 0.05. Top 1: study trait is the most diverged trait (with most “2” alleles) of *N* candidate loci. Random: study trait is chosen randomly from *N* candidate loci. Random50: study trait is chosen randomly from the 0.5 *N* candidate loci with the highest divergence (number of “2” alleles). Ratio: factor increase in false positive rate compared to observed rate when *N* = 1 for FPR from Random50.

|  | *N* | A&S Result | Top 1 | Random | Random50 | Ratio |
| --- | --- | --- | --- | --- | --- | --- |
| 11 Loci |  |  |  |  |  |  |
|  | TFPR | 0.0117 | 0.0117 | 0.0117 | 0.0117 |  |
|  | 1 | NA | 0.0120 | 0.0120 | 0.0120 | 1 |
|  | 10 | NA | 0.1153 | 0.0109 | 0.0243 | 2.025 |
|  | 25 | NA | 0.2622 | 0.0138 | 0.0224 | 1.867 |
|  | 50 | NA | 0.4505 | 0.013 | 0.0243 | 2.025 |
|  | 100 | NA | 0.6956 | 0.0124 | 0.0247 | 2.058 |
| 10 Loci | |  |  |  |  |  |
|  | TFPR | 0.0285 | 0.0285 | 0.0285 | 0.0285 |  |
|  | 1 | 0.0276 | 0.0290 | 0.0290 | 0.0290 | 1 |
|  | 10 | 0.2577 | 0.2518 | 0.0290 | 0.0587 | 2.024 |
|  | 25 | 0.5219 | 0.5144 | 0.0290 | 0.0519 | 1.790 |
|  | 50 | 0.7629 | 0.7647 | 0.0290 | 0.0568 | 1.959 |
|  | 100 | NA | 0.9477 | 0.0284 | 0.0579 | 1.997 |

Table A2. Ascertainment bias associated with the Fraser test. P value is calculated P value from Fraser test. Asterisk indicates P value remains significant at P < 0.05 after correction for ascertainment bias at a threshold P < 0.0075, corresponding to β > 0.15. Values in bold remain significant after a sequential Bonferroni correction. *N** is the maximum number of traits from which the indicated trait is selected (as the most diverged trait) in order for rejection of neutrality.

| Trait | P value | *N** |
| --- | --- | --- |
| Corolla Width | **0.00001526*** | 3,276 |
| Corolla Throat | **0.001*** | 50 |
| Corolla Length | **0.00001526*** | 3,276 |
| Overall Corolla Length | **0.00001526*** | 3,276 |
| Sepal Length | 0.49 | -- |
| Longest Stamen Length | **0.00001526*** | 3,276 |
| Pistil Length | 0.002 | 25 |
| Seed Mass | 0.35 | -- |
| Seed Length | 0.32 | -- |
| Seed Width | 0.32 | -- |
| Nectar Volume | **0.00001526*** | 3,276 |
| Nectar Sugar Concentration | **0.0001*** | 500 |
| Nectary Size | **0.0004*** | 125 |

Figure A1. False positive rate ratio (FPR Ratio) as a function of β. β is the proportion of 100 candidate traits with the highest divergence (highest number of “2” alleles) from which the study trait was chosen. β = 1 indicates study trait was chosen from all 100 candidate traits. β = 0.1 indicates study trait was chosen from the 10 percent of 100 traits with the highest divergence. FPR Ratio is the ratio (FPR for a given value of β)/(FPR for β = 1). Blue: simulated ratio. Red: expected ratio.

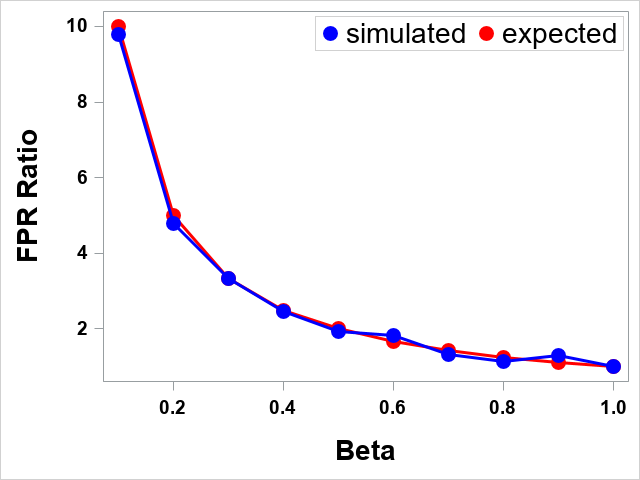

**Table S1** List of individuals phenotyped but NOT genotyped.

| F5 or Species | RIL (Line) | Individual |
| --- | --- | --- |
| F5 | 49 | 1272 |
| F5 | 88 | 1358 |
| F5 | 109 | 1151 |
| F5 | 142 | 1122 |
| F5 | 143 | 120 |
| F5 | 150 | 518 |
| F5 | 166 | 1338 |
| F5 | 177 | 923 |
| F5 | 250 | 895 |
| F5 | 291 | 88 |
| F5 | 295 | 5 |
| F5 | 331 | 695 |
| F5 | 344 | 178 |
| F5 | 375 | 165 |
| F5 | 411 | 706 |
| F5 | 438 | 1328 |
| F5 | 482 | 164 |
| F5 | 090b | 622 |
| F5 | 390b | 1321 |
| F5 | 474s | 1347 |
| F5 | 495s | 520 |
| L |  | 1 |
| L |  | 99 |

**Table S2** Genetic correlations derived from variance-covariance components for the 13 phenotypic traits measured. Genetic correlations are above the diagonal in the upper right triangle and environmental correlations are below the diagonal in the lower left triangle. Traits are in the same order as Figure 2. Each trait module is highlighted in gray.

|  | Seed Mass | Seed Length | Seed Width | Sugar Concentration | Nectar Volume | Nectary Size | Sepal Length | Corolla Width | Corolla Throat | Pistil Length | Longest Stamen Length | Corolla Length | Overall Corolla Length |
| --- | --- | --- | --- | --- | --- | --- | --- | --- | --- | --- | --- | --- | --- |
| Seed Mass | 1.000 | 0.702 | 0.714 | 0.046 | 0.060 | 0.148 | 0.271 | 0.145 | 0.166 | 0.100 | 0.217 | 0.185 | 0.143 |
| Seed Length |  | 1.000 | 0.719 | 0.126 | 0.103 | 0.263 | 0.391 | 0.232 | 0.288 | 0.291 | 0.281 | 0.212 | 0.199 |
| Seed Width |  |  | 1.000 | 0.144 | 0.045 | 0.123 | 0.178 | 0.162 | 0.127 | 0.116 | 0.232 | 0.188 | 0.158 |
| Sugar Concentration |  |  |  | 1.000 | 0.646 | 0.648 | 0.266 | 0.260 | 0.360 | 0.291 | 0.439 | 0.400 | 0.329 |
| Nectar Volume |  |  |  |  | 1.000 | 0.698 | 0.381 | 0.547 | 0.490 | 0.463 | 0.562 | 0.496 | 0.542 |
| Nectary Size |  |  |  |  |  | 1.000 | 0.465 | 0.545 | 0.607 | 0.395 | 0.446 | 0.412 | 0.445 |
| Sepal Length |  |  |  |  |  |  | 1.000 | 0.531 | 0.442 | 0.392 | 0.314 | 0.335 | 0.417 |
| Corolla Width |  |  |  |  |  |  |  | 1.000 | 0.698 | 0.588 | 0.672 | 0.635 | 0.787 |
| Corolla Throat |  |  |  |  |  |  |  |  | 1.000 | 0.409 | 0.440 | 0.543 | 0.589 |
| Pistil Length |  |  |  |  |  |  |  |  |  | 1.000 | 0.641 | 0.605 | 0.638 |
| Longest Stamen Length |  |  |  |  |  |  |  |  |  |  | 1.000 | 0.752 | 0.794 |
| Corolla Length |  |  |  |  |  |  |  |  |  |  |  | 1.000 | 0.978 |
| Overall Corolla Length |  |  |  |  |  |  |  |  |  |  |  |  | 1.000 |

**Table S3** QTL characteristics for all trait QTLs. a) Characteristics of GWS QTLs. b) Characteristics of CWS QTLs. L_M: QTL model peak derived from, L = *qtl2*, loco method; M = *qtl*, stepwiseqtl model (only for GWS QTLs). Chr: Chromosome. cM_pos: centimorgan position of the QTL peak. LOD: logarithm of the odds ratio of the QTL peak. CI_low: lower margin of 95% QTL confidence interval in centimorgans. CI_high: higher margin of the 95% QTL confidence interval in centimorgans. Direction: 1 = same direction as species difference; -1 = opposite direction. RHE: relative homozygous effect calculated from markers at the peak.

a) GWS (genome-wide significant) QTLs

| Trait | L_M | Chr | cM_pos | LOD | CI_low | CI_high | Direction | RHE |
| --- | --- | --- | --- | --- | --- | --- | --- | --- |
| CorollaWidth | M | 1 | 202.9555 | 9.341277 | 197.727 | 208.535 | 1 | 0.10847599 |
| CorollaWidth | M | 2 | 49.93117 | 8.669439 | 38.00915 | 55.90168 | 1 | 0.10248692 |
| CorollaWidth | M | 4 | 62.37565 | 6.470103 | 49.72713 | 79.26217 | 1 | 0.098879 |
| CorollaWidth | M | 5 | 167.7819 | 10.814064 | 163.7224 | 176.0234 | 1 | 0.1065059 |
| CorollaWidth | M | 8 | 227.0059 | 4.925853 | 210.8519 | 241.0999 | 1 | 0.06141165 |
| CorollaWidth | L | 10 | 177.9935 | 3.923731 | 144.6865 | 182.1695 | 1 | 0.08623802 |
| CorollaWidth | L | 11 | 111.0425 | 3.462357 | 100.2545 | 142.55 | 1 | 0.05040635 |
| CorollaWidth | M | 13 | 65.9782 | 10.362079 | 55.06614 | 74.0672 | 1 | 0.11162703 |
| CorollaThroat | M | 1 | 55.5166 | 4.358332 | 47.51458 | 70.90112 | 1 | 0.1102906 |
| CorollaThroat | M | 1 | 241.916 | 9.440827 | 226 | 248.8701 | 1 | 0.1716404 |
| CorollaThroat | M | 6 | 60.18907 | 3.998432 | 27.67103 | 65.35958 | 1 | 0.1342178 |
| CorollaThroat | M | 7 | 212 | 4.694686 | 201.7044 | 213.9574 | -1 | -0.1088578 |
| CorollaThroat | M | 8 | 192 | 3.143178 | 124.1033 | 208.7899 | 1 | 0.12414528 |
| CorollaThroat | M | 8 | 241.0999 | 5.694232 | 232.3009 | 241.0999 | 1 | 0.17609617 |
| CorollaThroat | L | 9 | 131.927 | 6.72557 | 103.85 | 147.6645 | 1 | 0.15341176 |
| CorollaThroat | L | 13 | 42.771 | 6.192714 | 34.8345 | 74.067 | 1 | 0.15362813 |
| CorollaLength | L | 1 | 207 | 5.488744 | 189.9165 | 231.566 | 1 | 0.08044557 |
| CorollaLength | M | 4 | 52 | 7.369602 | 45.27312 | 62.37565 | 1 | 0.10857611 |
| CorollaLength | M | 4 | 199.1425 | 3.362624 | 183.7614 | 227.0995 | 1 | 0.04722032 |
| CorollaLength | M | 5 | 70 | 7.596351 | 55.36365 | 87.19921 | 1 | 0.09750892 |
| CorollaLength | M | 5 | 171.0784 | 7.569567 | 163.7224 | 176.0234 | 1 | 0.11738669 |
| CorollaLength | M | 8 | 195.5423 | 5.915729 | 178.0508 | 204.6029 | 1 | 0.08830957 |
| CorollaLength | M | 9 | 27.4936 | 4.667363 | 0 | 38.07264 | 1 | 0.08714767 |
| CorollaLength | L | 9 | 145.207 | 13.851634 | 139.7295 | 147.6645 | 1 | 0.14135148 |
| CorollaLength | L | 14 | 138.9475 | 5.712576 | 129.887 | 143.2275 | -1 | -0.0711251 |
| CorollaLength | M | 15 | 27.21705 | 4.751402 | 6.18652 | 34.09405 | 1 | 0.10484593 |
| OverallCorollaLength | M | 1 | 204 | 8.781698 | 197.727 | 215.475 | 1 | 0.0860068 |
| OverallCorollaLength | L | 2 | 50 | 3.401029 | 0 | 56.5555 | 1 | 0.05330001 |
| OverallCorollaLength | M | 4 | 53 | 9.338872 | 46.09262 | 62.70165 | 1 | 0.09811914 |
| OverallCorollaLength | M | 5 | 74.93719 | 6.022362 | 59.71567 | 87.19921 | 1 | 0.0793927 |
| OverallCorollaLength | M | 5 | 171.0784 | 11.326458 | 163.7224 | 176.0234 | 1 | 0.12030212 |
| OverallCorollaLength | L | 8 | 199.001 | 5.762885 | 178.0505 | 215.694 | 1 | 0.06343492 |
| OverallCorollaLength | L | 9 | 145.207 | 12.213113 | 139.7295 | 147.6645 | 1 | 0.11464278 |
| OverallCorollaLength | L | 13 | 42.771 | 3.463942 | 32.847 | 74.067 | 1 | 0.06241793 |
| OverallCorollaLength | L | 14 | 138.9475 | 3.590595 | 127.2185 | 144.043 | -1 | -0.0473819 |
| SepalLength | M | 1 | 199 | 10.424359 | 195.1019 | 212.085 | 1 | 0.88956829 |
| SepalLength | M | 1 | 254 | 4.553586 | 240.196 | 267.2916 | 1 | 0.72622405 |
| SepalLength | M | 3 | 31.73156 | 3.957636 | 14.56553 | 39.26006 | -1 | -0.4706184 |
| SepalLength | M | 5 | 63.10468 | 4.10811827 | 2.373511 | 195.89843 | 1 | 0.39553961 |
| SepalLength | M | 5 | 196 | 5.938689 | 195 | 199 | 1 | 0.24649788 |
| SepalLength | M | 5 | 199 | 6.978751 | 196.5509 | 201.7014 | 1 | 0.35737443 |
| SepalLength | L | 6 | 201.194 | 4.226738 | 190.9315 | 227.2655 | -1 | -0.3803022 |
| SepalLength | M | 7 | 128.1678 | 3.630468 | 116.7277 | 135.1608 | -1 | -0.49571 |
| SepalLength | M | 9 | 38.07264 | 6.396238 | 32.49614 | 44.76115 | -1 | -0.4230288 |
| SepalLength | L | 12 | 144.152 | 5.89055 | 129.696 | 156.972 | -1 | -0.6099718 |
| SepalLength | M | 13 | 44.3356 | 3.846181 | 20.35354 | 74.0672 | 1 | 0.31943544 |
| SepalLength | M | 14 | 167.0247 | 6.668753 | 159.6697 | 181.2783 | 1 | 0.5787807 |
| LongestStamenLength | M | 1 | 5 | 3.453013 | 0 | 24.83956 | -1 | -0.0751853 |
| LongestStamenLength | M | 1 | 229.662 | 7.888189 | 222.529 | 240.196 | 1 | 0.09419313 |
| LongestStamenLength | M | 4 | 52 | 14.5691 | 46.09262 | 55.80463 | 1 | 0.12955287 |
| LongestStamenLength | L | 5 | 167.7815 | 5.405028 | 163.722 | 176.023 | 1 | 0.09532325 |
| LongestStamenLength | L | 8 | 201.2425 | 5.003138 | 178.0505 | 218.3195 | 1 | 0.05388055 |
| LongestStamenLength | M | 9 | 145.2074 | 8.112721 | 141.0459 | 147.6649 | 1 | 0.09854889 |
| LongestStamenLength | L | 11 | 27 | 5.76203 | 18.526 | 38.431 | 1 | 0.06759024 |
| LongestStamenLength | L | 13 | 58.3455 | 4.062428 | 19.7815 | 158.78 | 1 | 0.07173163 |
| LongestStamenLength | L | 14 | 141.24 | 6.878702 | 134.177 | 145.276 | -1 | -0.0749177 |
| PistilLength | M | 1 | 206 | 8.341122 | 195.1019 | 234.37 | 1 | 0.10079149 |
| PistilLength | M | 2 | 119.0478 | 3.677423 | 103.2068 | 130.7103 | -1 | -0.0784907 |
| PistilLength | M | 3 | 59 | 6.132963 | 49.21758 | 71.80162 | -1 | -0.1366132 |
| PistilLength | L | 4 | 52 | 20.080269 | 48.6605 | 64.3415 | 1 | 0.22534004 |
| PistilLength | M | 5 | 171.0784 | 15.60377 | 163.7224 | 176.0234 | 1 | 0.16584555 |
| PistilLength | M | 7 | 0 | 6.011584 | 0 | 23.56257 | 1 | 0.076754 |
| PistilLength | M | 7 | 142.4113 | 5.497754 | 128.1678 | 149.5473 | -1 | -0.0883203 |
| PistilLength | L | 7 | 192.457 | 5.401624 | 184.371 | 198.645 | -1 | -0.0873558 |
| PistilLength | M | 8 | 64.22617 | 4.382283 | 55.06363 | 84.46073 | -1 | -0.0084587 |
| PistilLength | M | 8 | 214 | 8.714308 | 199.0014 | 227.0059 | 1 | 0.12690483 |
| PistilLength | M | 9 | 12.70103 | 17.77145 | 9.603527 | 22.733586 | 1 | 0.17724894 |
| PistilLength | M | 9 | 135.4704 | 3.947435 | 102 | 147.6649 | 1 | 0.10092129 |
| PistilLength | M | 10 | 58.91763 | 13.00417 | 55.09462 | 61.84413 | 1 | 0.06301991 |
| PistilLength | M | 13 | 51 | 5.526912 | 39.32659 | 61.86767 | 1 | 0.11535459 |
| PistilLength | M | 13 | 151.3448 | 8.080634 | 149.9528 | 160.9903 | 1 | 0.1051753 |
| PistilLength | M | 15 | 9 | 3.480625 | 0 | 34.09405 | 1 | 0.09193956 |
| SeedMass | M | 12 | 146.1997 | 3.328422 | 100.5231 | 156.9722 | 1 | 0.37459617 |
| SeedLength | L | 2 | 92.3455 | 3.196395 | 13.49 | 107.2625 | 1 | 0.30016197 |
| SeedLength | M | 12 | 115.7787 | 3.773297 | 92.62264 | 132.79419 | 1 | 0.35787316 |
| SeedLength | M | 13 | 65.9782 | 5.146565 | 15.32403 | 74.0672 | -1 | -0.4262481 |
| SeedWidth | M | 1 | 192.6404 | 3.830288 | 150.6488 | 226.785 | -1 | -0.371878 |
| NectarVolume | L | 4 | 154.002 | 3.656793 | 121.5925 | 171.8655 | 1 | 0.07200126 |
| NectarVolume | L | 5 | 26 | 3.229344 | 2.3735 | 69.7335 | 1 | 0.07904937 |
| NectarVolume | L | 13 | 69 | 3.807356 | 25.0625 | 145.988 | 1 | 0.09606644 |
| SugarConcentration | M | 4 | 197.1545 | 8.587753 | 183.7614 | 205.542 | 1 | 0.12176364 |
| SugarConcentration | M | 7 | 84.79965 | 3.473199 | 36.74659 | 94.4942 | 1 | 0.09115739 |
| SugarConcentration | M | 9 | 137 | 5.078752 | 59.05767 | 146.93187 | 1 | 0.10327198 |
| SugarConcentration | M | 10 | 72.60815 | 4.340746 | 65.72614 | 94.17273 | 1 | 0.09068293 |
| SugarConcentration | M | 11 | 25.88959 | 5.917689 | 18.52608 | 34.44509 | 1 | 0.12862033 |
| SugarConcentration | M | 15 | 60.17856 | 3.824674 | 56.21056 | 74.61958 | 1 | 0.11824147 |
| NectarySize | M | 1 | 203 | 5.349908 | 189.9169 | 232.2205 | 1 | 0.09675231 |
| NectarySize | M | 3 | 51 | 8.234195 | 47.16808 | 55.91109 | -1 | -0.2080672 |
| NectarySize | L | 4 | 186 | 6.532344 | 171.8655 | 196.253 | 1 | 0.10801165 |
| NectarySize | L | 5 | 52.4275 | 5.665011 | 45.308 | 66.2895 | 1 | 0.10505225 |
| NectarySize | M | 6 | 167 | 3.676935 | 153.5923 | 195.2539 | 1 | 0.07264236 |
| NectarySize | M | 7 | 205.7089 | 5.351719 | 196.2504 | 212.4894 | -1 | -0.1000032 |
| NectarySize | M | 8 | 30.58759 | 7.936686 | 24.01307 | 44.75011 | -1 | -0.1136726 |
| NectarySize | M | 13 | 0.244014 | 7.314697 | 0 | 28.767083 | 1 | 0.10123476 |
| NectarySize | M | 13 | 142.9588 | 5.083236 | 134 | 149.7898 | 1 | 0.10379771 |
| NectarySize | M | 14 | 20 | 8.008347 | 12 | 29 | 1 | 0.12231527 |
| NectarySize | M | 14 | 165.8767 | 7.151813 | 155.4762 | 185.4798 | 1 | 0.11881932 |

b) CWS (chromosome-wide significant) QTLs

| Trait | Chr | cM_Pos | LOD | ci_lo | ci_hi | RHE | Direction |
| --- | --- | --- | --- | --- | --- | --- | --- |
| CorollaWidth | 1 | 69 | 2.931533 | 52.277 | 83.172 | 0.08078033 | 1 |
| CorollaWidth | 1 | 205 | 10.140215 | 196.4175 | 231.566 | 0.10761747 | 1 |
| CorollaWidth | 2 | 49.931 | 8.753845 | 38.009 | 55.9015 | 0.10248692 | 1 |
| CorollaWidth | 3 | 141 | 3.097203 | 84.789 | 167.9645 | 0.03590189 | 1 |
| CorollaWidth | 4 | 62.3755 | 6.171439 | 46.0925 | 96.087 | 0.098879 | 1 |
| CorollaWidth | 5 | 167.7815 | 7.534143 | 153.3715 | 176.023 | 0.1065059 | 1 |
| CorollaWidth | 5 | 201.701 | 7.23343 | 196.5505 | 204.4765 | 0.0990126 | 1 |
| CorollaWidth | 8 | 206.8125 | 6.093384 | 178.0505 | 232.3005 | 0.05604645 | 1 |
| CorollaWidth | 9 | 145.207 | 2.564658 | 103.85 | 147.6645 | 0.0624393 | 1 |
| CorollaWidth | 10 | 177.9935 | 3.923731 | 144.6865 | 182.1695 | 0.08623802 | 1 |
| CorollaWidth | 11 | 111.0425 | 3.462357 | 100.2545 | 142.55 | 0.05040635 | 1 |
| CorollaWidth | 13 | 65.978 | 8.789049 | 55.066 | 74.067 | 0.11162703 | 1 |
| CorollaWidth | 14 | 97 | 2.415352 | 0 | 111.1115 | 0.05010576 | 1 |
| CorollaThroat | 1 | 61.731 | 4.446111 | 47.5145 | 80.3005 | 0.1109311 | 1 |
| CorollaThroat | 1 | 242.2405 | 9.844063 | 227.2735 | 254.236 | 0.17166973 | 1 |
| CorollaThroat | 3 | 52 | 2.456232 | 39.9135 | 212.4895 | -0.1521561 | -1 |
| CorollaThroat | 4 | 224 | 2.259363 | 180.8085 | 227.099 | 0.09172691 | 1 |
| CorollaThroat | 5 | 201.211 | 2.136158 | 128.866 | 204.4765 | 0.05263552 | 1 |
| CorollaThroat | 6 | 57 | 2.89354 | 11.937 | 227.2655 | 0.12727942 | 1 |
| CorollaThroat | 7 | 211.835 | 5.747755 | 201.704 | 213.957 | -0.1086791 | -1 |
| CorollaThroat | 8 | 227.0865 | 9.752778 | 212.4055 | 232.3005 | 0.17562306 | 1 |
| CorollaThroat | 8 | 241.0185 | 9.624274 | 232.3005 | 241.0995 | 0.17656928 | 1 |
| CorollaThroat | 9 | 131.927 | 6.72557 | 103.85 | 147.6645 | 0.15341176 | 1 |
| CorollaThroat | 12 | 148.696 | 2.11408 | 134.9235 | 185.037 | -0.0405575 | -1 |
| CorollaThroat | 13 | 42.771 | 6.192714 | 34.8345 | 74.067 | 0.1391421 | 1 |
| CorollaThroat | 14 | 2.126 | 2.149904 | 0 | 60.1655 | 0.07531132 | 1 |
| CorollaThroat | 15 | 10.0715 | 3.010917 | 0 | 34.094 | 0.08075448 | 1 |
| CorollaLength | 1 | 207 | 5.488744 | 189.9165 | 231.566 | 0.08476221 | 1 |
| CorollaLength | 4 | 52 | 8.974564 | 36.826 | 64.3415 | 0.10857611 | 1 |
| CorollaLength | 4 | 199.142 | 3.421336 | 151.503 | 220.97 | 0.04722032 | 1 |
| CorollaLength | 5 | 69.7335 | 7.11869 | 55.3635 | 103.139 | 0.09694614 | 1 |
| CorollaLength | 5 | 169 | 7.902297 | 156.1125 | 176.023 | 0.11701555 | 1 |
| CorollaLength | 6 | 164.3355 | 2.381859 | 0 | 227.2655 | -0.0438315 | -1 |
| CorollaLength | 7 | 25.6185 | 2.138707 | 0 | 110.95 | 0.02874012 | 1 |
| CorollaLength | 8 | 195.216 | 5.748446 | 178.0505 | 215.694 | 0.08740428 | 1 |
| CorollaLength | 9 | 35 | 5.64683 | 26.35 | 52.792 | 0.09422588 | 1 |
| CorollaLength | 9 | 145.207 | 13.851634 | 139.7295 | 147.6645 | 0.1407039 | 1 |
| CorollaLength | 13 | 42.771 | 1.866423 | 0 | 74.067 | 0.06078641 | 1 |
| CorollaLength | 14 | 138.9475 | 5.712576 | 129.887 | 143.2275 | -0.0714593 | -1 |
| CorollaLength | 15 | 27.217 | 3.806571 | 4.03 | 34.094 | 0.10484593 | 1 |
| OverallCorollaLength | 1 | 207 | 8.237548 | 197.6455 | 224.9825 | 0.08712264 | 1 |
| OverallCorollaLength | 2 | 50 | 3.401029 | 0 | 56.5555 | 0.05330001 | 1 |
| OverallCorollaLength | 3 | 151.9625 | 2.418239 | 88.778 | 187.352 | 0.04741774 | 1 |
| OverallCorollaLength | 4 | 52 | 9.179843 | 45.273 | 64.3415 | 0.09811914 | 1 |
| OverallCorollaLength | 4 | 199.142 | 2.874222 | 145.9415 | 220.97 | 0.03753586 | 1 |
| OverallCorollaLength | 5 | 98 | 6.337307 | 60.782 | 105.184 | 0.07428009 | 1 |
| OverallCorollaLength | 5 | 169 | 11.680557 | 163.722 | 176.023 | 0.11998197 | 1 |
| OverallCorollaLength | 5 | 202 | 7.90442 | 196.5505 | 204.4765 | 0.09543185 | 1 |
| OverallCorollaLength | 7 | 25.6185 | 2.633428 | 0 | 89.655 | 0.03089939 | 1 |
| OverallCorollaLength | 8 | 199.001 | 5.762885 | 178.0505 | 215.694 | 0.06216653 | 1 |
| OverallCorollaLength | 9 | 28 | 5.042618 | 22.7335 | 46.476 | 0.06732063 | 1 |
| OverallCorollaLength | 9 | 145.207 | 12.213113 | 139.7295 | 147.6645 | 0.11514436 | 1 |
| OverallCorollaLength | 13 | 42.771 | 3.463942 | 32.847 | 74.067 | 0.06241793 | 1 |
| OverallCorollaLength | 14 | 138.9475 | 3.590595 | 127.2185 | 144.043 | -0.0473819 | -1 |
| SepalLength | 1 | 70 | 4.417509 | 58.4255 | 138.0135 | 0.55613191 | 1 |
| SepalLength | 1 | 206 | 13.015013 | 196.4175 | 214.2415 | 0.89687411 | 1 |
| SepalLength | 4 | 35.924 | 2.040663 | 10.419 | 171.8655 | 0.33971682 | 1 |
| SepalLength | 5 | 69.7335 | 3.060204 | 0 | 203.661 | 0.37577092 | 1 |
| SepalLength | 6 | 201.194 | 4.226738 | 190.9315 | 227.2655 | -0.3727099 | -1 |
| SepalLength | 7 | 128.1675 | 3.405859 | 84.7995 | 201.4605 | -0.49571 | -1 |
| SepalLength | 9 | 38.0725 | 5.129109 | 26.35 | 44.761 | -0.4230288 | -1 |
| SepalLength | 11 | 190.251 | 2.517069 | 6.241 | 190.251 | -0.2597976 | -1 |
| SepalLength | 12 | 144.152 | 5.89055 | 129.696 | 156.972 | -0.5949304 | -1 |
| SepalLength | 14 | 180.5415 | 6.645739 | 159.6695 | 185.4795 | 0.60860714 | 1 |
| SepalLength | 15 | 0 | 1.925394 | 0 | 129.8885 | 0.31786721 | 1 |
| LongestStamenLength | 1 | 6.171 | 2.559498 | 0 | 22.852 | -0.074194 | -1 |
| LongestStamenLength | 1 | 229.008 | 6.873327 | 202.955 | 237.407 | 0.09424421 | 1 |
| LongestStamenLength | 2 | 30.8095 | 3.275676 | 2.616 | 56.5555 | 0.04837596 | 1 |
| LongestStamenLength | 4 | 46.0925 | 13.933133 | 36.826 | 64.3415 | 0.12591072 | 1 |
| LongestStamenLength | 4 | 183.761 | 3.251964 | 151.503 | 218.1885 | 0.04948091 | 1 |
| LongestStamenLength | 5 | 89.571 | 3.435663 | 59.7155 | 102.318 | 0.05924846 | 1 |
| LongestStamenLength | 5 | 167.7815 | 5.405028 | 163.722 | 176.023 | 0.09532326 | 1 |
| LongestStamenLength | 7 | 43 | 2.906178 | 0 | 211.347 | 0.03563166 | 1 |
| LongestStamenLength | 8 | 201.2425 | 5.003138 | 178.0505 | 218.3195 | 0.05388055 | 1 |
| LongestStamenLength | 9 | 145.207 | 7.196786 | 135.47 | 147.6645 | 0.09854889 | 1 |
| LongestStamenLength | 11 | 27 | 5.76203 | 18.526 | 38.431 | 0.06759024 | 1 |
| LongestStamenLength | 13 | 58.3455 | 4.062428 | 19.7815 | 74.067 | 0.07108492 | 1 |
| LongestStamenLength | 13 | 142.9585 | 3.885541 | 132.6865 | 158.78 | 0.0711065 | 1 |
| LongestStamenLength | 14 | 141.24 | 6.878702 | 134.177 | 145.276 | -0.0749177 | -1 |
| PistilLength | 1 | 224.249 | 8.957204 | 201.805 | 244.152 | 0.10201085 | 1 |
| PistilLength | 2 | 112.3145 | 4.873913 | 56.5555 | 130.71 | -0.0823761 | -1 |
| PistilLength | 3 | 64.0185 | 3.234325 | 26.7175 | 78.8795 | -0.1282076 | -1 |
| PistilLength | 4 | 52 | 20.080269 | 48.6605 | 57.7715 | 0.22895452 | 1 |
| PistilLength | 4 | 61 | 19.776475 | 57.7715 | 69.3825 | 0.22172555 | 1 |
| PistilLength | 4 | 182.9425 | 2.8457 | 171.8655 | 224.7925 | 0.0712001 | 1 |
| PistilLength | 5 | 167.7815 | 9.487924 | 163.722 | 176.023 | 0.16944393 | 1 |
| PistilLength | 5 | 201.701 | 6.624499 | 196.5505 | 204.4765 | 0.13503102 | 1 |
| PistilLength | 7 | 0 | 3.134612 | 0 | 23.5625 | 0.076754 | 1 |
| PistilLength | 7 | 192.457 | 5.401624 | 184.371 | 198.645 | -0.0873558 | -1 |
| PistilLength | 8 | 30.5875 | 2.583274 | 23.2775 | 91.61 | -0.0299656 | -1 |
| PistilLength | 8 | 215 | 6.887641 | 178.0505 | 223.1235 | 0.12881392 | 1 |
| PistilLength | 9 | 12.701 | 14.076313 | 9.6035 | 31.76 | 0.17724894 | 1 |
| PistilLength | 9 | 145.207 | 4.347136 | 132.4975 | 147.6645 | 0.10955815 | 1 |
| PistilLength | 10 | 58.9175 | 5.117359 | 42.8065 | 65.726 | 0.06301991 | 1 |
| PistilLength | 13 | 46.0535 | 5.342284 | 39.3265 | 74.067 | 0.11352224 | 1 |
| PistilLength | 13 | 142.9585 | 8.335773 | 132.6865 | 158.78 | 0.10055558 | 1 |
| PistilLength | 14 | 96 | 2.209339 | 65.4305 | 183.012 | 0.0982173 | 1 |
| PistilLength | 15 | 10 | 2.85403 | 0 | 34.094 | 0.09425889 | 1 |
| SeedMass | 1 | 193.618 | 2.269288 | 150.6485 | 265.98 | -0.304447 | -1 |
| SeedMass | 12 | 133.2835 | 2.835046 | 100.523 | 156.972 | 0.35804688 | 1 |
| SeedMass | 13 | 65 | 1.881849 | 15.324 | 160.99 | -0.3051093 | -1 |
| SeedLength | 1 | 260.3905 | 2.402093 | 1.628 | 268.514 | -0.2419011 | -1 |
| SeedLength | 2 | 92.3455 | 3.196395 | 13.49 | 107.2625 | 0.30016197 | 1 |
| SeedLength | 4 | 218.1885 | 2.299019 | 0 | 227.099 | -0.3176993 | -1 |
| SeedLength | 11 | 16.483 | 2.599628 | 0.6515 | 115.595 | -0.3681337 | -1 |
| SeedLength | 12 | 116 | 3.302696 | 100.523 | 141.3395 | 0.36056283 | 1 |
| SeedLength | 13 | 24.9815 | 5.34003 | 15.324 | 74.067 | -0.4236709 | -1 |
| SeedLength | 14 | 182 | 3.104524 | 163.464 | 185.4795 | -0.2960413 | -1 |
| SeedWidth | 1 | 193.618 | 3.569231 | 150.2405 | 238.392 | -0.376416 | -1 |
| SeedWidth | 12 | 119 | 3.047802 | 100.523 | 148.696 | 0.36858483 | 1 |
| NectarVolume | 1 | 191.904 | 2.293941 | 10.6385 | 212.0845 | 0.0635895 | 1 |
| NectarVolume | 3 | 196.4575 | 2.631956 | 49.2175 | 212.4895 | 0.08611989 | 1 |
| NectarVolume | 4 | 154.002 | 3.656793 | 121.5925 | 171.8655 | 0.07200126 | 1 |
| NectarVolume | 5 | 26 | 3.229344 | 2.3735 | 69.7335 | 0.07904937 | 1 |
| NectarVolume | 7 | 205.7085 | 2.337015 | 24.0515 | 213.957 | -0.0795507 | -1 |
| NectarVolume | 8 | 216.4295 | 2.075774 | 0 | 241.0995 | 0.07496359 | 1 |
| NectarVolume | 10 | 2.124 | 2.64068 | 0 | 27.415 | 0.07455409 | 1 |
| NectarVolume | 13 | 69 | 3.807356 | 25.0625 | 85.278 | 0.09682594 | 1 |
| NectarVolume | 13 | 142.9585 | 3.426826 | 107.6085 | 151.019 | 0.0954645 | 1 |
| SugarConcentration | 4 | 197 | 6.595214 | 180.8085 | 218.1885 | 0.12194979 | 1 |
| SugarConcentration | 5 | 23.6985 | 2.201672 | 4.0915 | 66.2895 | 0.08335363 | 1 |
| SugarConcentration | 7 | 84.7995 | 2.890754 | 24.0515 | 105.2735 | 0.09115739 | 1 |
| SugarConcentration | 8 | 31.3235 | 2.486823 | 0 | 241.0995 | -0.077572 | -1 |
| SugarConcentration | 9 | 68.0645 | 4.118054 | 57.908 | 78.6575 | 0.08453126 | 1 |
| SugarConcentration | 9 | 136.3695 | 4.801412 | 132.4975 | 146.9315 | 0.1051972 | 1 |
| SugarConcentration | 10 | 77.4425 | 2.653526 | 40.138 | 100.9855 | 0.0880649 | 1 |
| SugarConcentration | 11 | 25.8895 | 5.72575 | 18.526 | 45.788 | 0.12862033 | 1 |
| SugarConcentration | 13 | 6 | 2.271226 | 0 | 151.9985 | 0.06132722 | 1 |
| SugarConcentration | 14 | 163 | 2.363912 | 138.9475 | 185.4795 | 0.09872577 | 1 |
| SugarConcentration | 15 | 60.1785 | 3.267661 | 56.2105 | 74.6195 | 0.11824147 | 1 |
| NectarySize | 1 | 70.901 | 2.633468 | 52.277 | 142.6955 | 0.07840705 | 1 |
| NectarySize | 1 | 204 | 4.030496 | 181.6895 | 241.0965 | 0.09319656 | 1 |
| NectarySize | 3 | 51 | 6.986244 | 45.6875 | 55.911 | -0.2080672 | -1 |
| NectarySize | 4 | 186 | 6.532344 | 171.8655 | 196.253 | 0.11083411 | 1 |
| NectarySize | 5 | 52.4275 | 5.665011 | 45.308 | 66.2895 | 0.10006939 | 1 |
| NectarySize | 6 | 159 | 3.117219 | 128.4285 | 190.3605 | 0.0890567 | 1 |
| NectarySize | 7 | 205.627 | 3.606387 | 196.25 | 213.957 | -0.1000032 | -1 |
| NectarySize | 8 | 30.5875 | 6.007034 | 27.4625 | 51.358 | -0.1136726 | -1 |
| NectarySize | 8 | 227.0865 | 3.190246 | 205.669 | 241.0995 | 0.0730956 | 1 |
| NectarySize | 11 | 158.611 | 2.438949 | 18.526 | 187.886 | 0.0623858 | 1 |
| NectarySize | 12 | 172.178 | 2.457566 | 141.584 | 193.7315 | -0.0505758 | -1 |
| NectarySize | 13 | 24.9815 | 7.947556 | 10.7365 | 32.847 | 0.12854911 | 1 |
| NectarySize | 13 | 58.3455 | 8.107684 | 34.8345 | 77.65 | 0.13569644 | 1 |
| NectarySize | 13 | 142 | 5.616595 | 107.6085 | 146.6415 | 0.10115525 | 1 |
| NectarySize | 14 | 18 | 7.583916 | 6.173 | 32.1965 | 0.12916037 | 1 |
| NectarySize | 14 | 184 | 6.311883 | 153.596 | 185.4795 | 0.12521122 | 1 |

**Table S4** Summary of aggregate QTL characteristics for individual traits. a) Characteristics of GWS QTLs. b) Characteristics of ALL QTLs. TRHE: Total relative homozygous effect (RHE) of all QTLs, including those with both positive and negative effects. Mean RHE: Mean of absolute values of RHE. Max RHE: maximum of absolute values of RHE. CV RHE: Coefficient of Variation of absolute values of RHE. Consistent Direction: Proportion of QTLs with effects in same direction as the difference between species (Total number of QTLs). Test P: Probability of QTL-EE sign test under the null hypothesis of no selection. Bold values indicate significance at an overall level of P < 0.05 after a Bonferroni correction for multiple comparisons. NA: Test not conducted because number of QTLs less than 8. Corrected Test P: significance level adjusted for ascertainment bias assuming β > 0.15 (see text).

| Trait | Module | TRHE | Mean | Max | CV | Consistent | Test P | Corrected Test P |
| --- | --- | --- | --- | --- | --- | --- | --- | --- |
|  |  |  | RHE | RHE* | RHE* | Direction |  |  |
| Corolla Width | flower | 0.726 | 0.091 | 0.112 | 0.2537 | 1.000 (8) | **0.00391** | 0.026 |
| Corolla Throat | flower | 0.915 | 0.114 | 0.176 | 0.1847 | 0.875 (8) | 0.03135 | 0.210 |
| Corolla Length | flower | 0.802 | 0.08 | 0.141 | 0.2759 | 0.900 (10) | **0.00977** | 0.065 |
| Overall Corolla Length | flower | 0.63 | 0.07 | 0.12 | 0.3268 | 0.888 (9) | 0.01758 | 0.119 |
| Sepal Length | flower | 1.134 | 0.094 | 0.89 | 0.3732 | 0.583 (12) | 0.19336 | 1.00 |
| Longest Stamen Length | flower | 0.461 | 0.051 | 0.13 | 0.2646 | 0.778 (9) | 0.07031 | 0.469 |
| Pistil Length | flower | 0.95 | 0.059 | 0.225 | 0.461 | 0.688 (16) | 0.06665 | 0.444 |
| Seed Mass | seed | 0.375 | 0.375 | 0.375 | NA | 1.000 (1) | NA | NA |
| Seed Length | seed | 0.232 | 0.077 | 0.426 | 0.1746 | 0.667 (3) | NA | NA |
| Seed Width | seed | -0.372 | -0.372 | 0.372 | NA | 0.000 (1) | NA | NA |
| Nectar Volume | nectar | 0.247 | 0.082 | 0.096 | 0.1502 | 1.000 (3) | NA | NA |
| Nectar Sugar Concentration | nectar | 0.654 | 0.109 | 0.129 | 0.1491 | 1.000 (6) | NA | NA |
| Nectary Size | nectar | 0.407 | 0.037 | 0.208 | 0.2987 | 0.727 (11) | 0.08057 | 0.537 |

a)

b)

| Trait | Module | TRHE | Mean | Max | CV | Consistent | Test P | Corrected Test P |
| --- | --- | --- | --- | --- | --- | --- | --- | --- |
|  |  |  | RHE | RHE* | RHE* | Direction |  |  |
| Corolla Width | flower | 1.048 | 0.081 | 0.112 | 0.327 | 1.0 (13) | **0.000122** | 0.000813 |
| Corolla Throat | flower | 1.054 | 0.075 | 0.177 | 0.384 | 0.786 (14) | 0.0222 | 0.148 |
| Corolla Length | flower | 0.856 | 0.066 | 0.141 | 0.384 | 0.846 (13) | **0.0095** | 0.063 |
| Overall Corolla Length | flower | 0.904 | 0.065 | 0.12 | 0.394 | 0.929 (14) | **0.000854** | 0.00569 |
| Sepal Length | flower | 0.949 | 0.086 | 0.897 | 0.381 | 0.545 (11) | 0.2256 | 1.00 |
| Longest Stamen Length | flower | 0.721 | 0.052 | 0.126 | 0.331 | 0.857 (14) | **0.0056** | 0.0373 |
| Pistil Length | flower | 1.562 | 0.082 | 0.229 | 0.443 | 0.790 (19) | **0.00739** | 0.0490 |
| Seed Mass | seed | -0.252 | -0.084 | 0.358 | 0.095 | 0.333 (3) | NA | -- |
| Seed Length | seed | -0.987 | -0.141 | 0.423 | 0.18 | 0.286 (7) | NA | -- |
| Seed Width | seed | -0.008 | -0.004 | 0.376 | 0.015 | 0.5 (2) | NA | -- |
| Nectar Volume | nectar | 0.563 | 0.063 | 0.097 | 0.136 | 0.889 (9) | 0.0176 | 0.117 |
| Nectar Sugar Concentration | nectar | 0.904 | 0.082 | 0.129 | 0.214 | 0.909 (11) | **0.00537** | 0.0358 |
| Nectary Size | nectar | 0.754 | 0.047 | 0.208 | 0.345 | 0.750 (16) | 0.02778 | 0.185 |

**Table S5** Pairwise QTL overlap. a) List based on GWS (genome-wide significant) QTLs. b) List based on ALL QTLs. nQTL1: number of QTLs for trait 1. nQTL2: number of QTLs for trait 2. nOverlap: overlapping QTLs between traits. Jaccard_Overlap: Jaccard index of QTL overlap for the two traits.

a) GWS QTLs

| Pairwise Traits | nQTL1 | nQTL2 | nOverlap | Jaccard_Overlap |
| --- | --- | --- | --- | --- |
| CorollaLength-CorollaLength | 10 | 10 | 10 | 1 |
| CorollaLength-CorollaThroat | 10 | 8 | 3 | 0.2 |
| CorollaLength-CorollaWidth | 10 | 8 | 3 | 0.2 |
| CorollaLength-LongestStamenLength | 10 | 9 | 6 | 0.46153846 |
| CorollaLength-NectarVolume | 10 | 3 | 1 | 0.08333333 |
| CorollaLength-NectarySize | 10 | 11 | 3 | 0.16666667 |
| CorollaLength-OverallCorollaLength | 10 | 9 | 7 | 0.58333333 |
| CorollaLength-PistilLength | 10 | 16 | 7 | 0.36842105 |
| CorollaLength-SeedLength | 10 | 3 | 0 | 0 |
| CorollaLength-SeedMass | 10 | 1 | 0 | 0 |
| CorollaLength-SeedWidth | 10 | 1 | 1 | 0.1 |
| CorollaLength-SepalLength | 10 | 12 | 4 | 0.22222222 |
| CorollaLength-SugarConcentration | 10 | 6 | 2 | 0.14285714 |
| CorollaThroat-CorollaThroat | 8 | 8 | 8 | 1 |
| CorollaThroat-CorollaWidth | 8 | 8 | 2 | 0.14285714 |
| CorollaThroat-LongestStamenLength | 8 | 9 | 4 | 0.30769231 |
| CorollaThroat-NectarVolume | 8 | 3 | 1 | 0.1 |
| CorollaThroat-NectarySize | 8 | 11 | 2 | 0.11764706 |
| CorollaThroat-OverallCorollaLength | 8 | 9 | 3 | 0.21428571 |
| CorollaThroat-PistilLength | 8 | 16 | 4 | 0.2 |
| CorollaThroat-SeedLength | 8 | 3 | 1 | 0.1 |
| CorollaThroat-SeedMass | 8 | 1 | 0 | 0 |
| CorollaThroat-SeedWidth | 8 | 1 | 1 | 0.125 |
| CorollaThroat-SepalLength | 8 | 12 | 2 | 0.11111111 |
| CorollaThroat-SugarConcentration | 8 | 6 | 1 | 0.07692308 |
| CorollaWidth-CorollaWidth | 8 | 8 | 8 | 1 |
| CorollaWidth-LongestStamenLength | 8 | 9 | 4 | 0.30769231 |
| CorollaWidth-NectarVolume | 8 | 3 | 1 | 0.1 |
| CorollaWidth-NectarySize | 8 | 11 | 1 | 0.05555556 |
| CorollaWidth-OverallCorollaLength | 8 | 9 | 6 | 0.54545455 |
| CorollaWidth-PistilLength | 8 | 16 | 5 | 0.26315789 |
| CorollaWidth-SeedLength | 8 | 3 | 2 | 0.22222222 |
| CorollaWidth-SeedMass | 8 | 1 | 0 | 0 |
| CorollaWidth-SeedWidth | 8 | 1 | 1 | 0.125 |
| CorollaWidth-SepalLength | 8 | 12 | 3 | 0.17647059 |
| CorollaWidth-SugarConcentration | 8 | 6 | 0 | 0 |
| LongestStamenLength-LongestStamenLength | 9 | 9 | 9 | 1 |
| LongestStamenLength-NectarVolume | 9 | 3 | 1 | 0.09090909 |
| LongestStamenLength-NectarySize | 9 | 11 | 3 | 0.17647059 |
| LongestStamenLength-OverallCorollaLength | 9 | 9 | 6 | 0.5 |
| LongestStamenLength-PistilLength | 9 | 16 | 7 | 0.38888889 |
| LongestStamenLength-SeedLength | 9 | 3 | 1 | 0.09090909 |
| LongestStamenLength-SeedMass | 9 | 1 | 0 | 0 |
| LongestStamenLength-SeedWidth | 9 | 1 | 1 | 0.11111111 |
| LongestStamenLength-SepalLength | 9 | 12 | 2 | 0.10526316 |
| LongestStamenLength-SugarConcentration | 9 | 6 | 2 | 0.15384615 |
| NectarVolume-NectarVolume | 3 | 3 | 3 | 1 |
| NectarVolume-NectarySize | 3 | 11 | 3 | 0.27272727 |
| NectarVolume-OverallCorollaLength | 3 | 9 | 2 | 0.2 |
| NectarVolume-PistilLength | 3 | 16 | 1 | 0.05555556 |
| NectarVolume-SeedLength | 3 | 3 | 1 | 0.2 |
| NectarVolume-SeedMass | 3 | 1 | 0 | 0 |
| NectarVolume-SeedWidth | 3 | 1 | 0 | 0 |
| NectarVolume-SepalLength | 3 | 12 | 2 | 0.15384615 |
| NectarVolume-SugarConcentration | 3 | 6 | 0 | 0 |
| NectarySize-NectarySize | 11 | 11 | 11 | 1 |
| NectarySize-OverallCorollaLength | 11 | 9 | 2 | 0.11111111 |
| NectarySize-PistilLength | 11 | 16 | 3 | 0.125 |
| NectarySize-SeedLength | 11 | 3 | 1 | 0.07692308 |
| NectarySize-SeedMass | 11 | 1 | 0 | 0 |
| NectarySize-SeedWidth | 11 | 1 | 1 | 0.09090909 |
| NectarySize-SepalLength | 11 | 12 | 5 | 0.27777778 |
| NectarySize-SugarConcentration | 11 | 6 | 1 | 0.0625 |
| OverallCorollaLength-OverallCorollaLength | 9 | 9 | 9 | 1 |
| OverallCorollaLength-PistilLength | 9 | 16 | 6 | 0.31578947 |
| OverallCorollaLength-SeedLength | 9 | 3 | 2 | 0.2 |
| OverallCorollaLength-SeedMass | 9 | 1 | 0 | 0 |
| OverallCorollaLength-SeedWidth | 9 | 1 | 1 | 0.11111111 |
| OverallCorollaLength-SepalLength | 9 | 12 | 4 | 0.23529412 |
| OverallCorollaLength-SugarConcentration | 9 | 6 | 1 | 0.07142857 |
| PistilLength-PistilLength | 16 | 16 | 16 | 1 |
| PistilLength-SeedLength | 16 | 3 | 2 | 0.11764706 |
| PistilLength-SeedMass | 16 | 1 | 0 | 0 |
| PistilLength-SeedWidth | 16 | 1 | 1 | 0.0625 |
| PistilLength-SepalLength | 16 | 12 | 4 | 0.16666667 |
| PistilLength-SugarConcentration | 16 | 6 | 1 | 0.04761905 |
| SeedLength-SeedLength | 3 | 3 | 3 | 1 |
| SeedLength-SeedMass | 3 | 1 | 1 | 0.33333333 |
| SeedLength-SeedWidth | 3 | 1 | 0 | 0 |
| SeedLength-SepalLength | 3 | 12 | 2 | 0.15384615 |
| SeedLength-SugarConcentration | 3 | 6 | 0 | 0 |
| SeedMass-SeedMass | 1 | 1 | 1 | 1 |
| SeedMass-SeedWidth | 1 | 1 | 0 | 0 |
| SeedMass-SepalLength | 1 | 12 | 1 | 0.08333333 |
| SeedMass-SugarConcentration | 1 | 6 | 0 | 0 |
| SeedWidth-SeedWidth | 1 | 1 | 1 | 1 |
| SeedWidth-SepalLength | 1 | 12 | 1 | 0.08333333 |
| SeedWidth-SugarConcentration | 1 | 6 | 0 | 0 |
| SepalLength-SepalLength | 12 | 12 | 12 | 1 |
| SepalLength-SugarConcentration | 12 | 6 | 0 | 0 |
| SugarConcentration-SugarConcentration | 6 | 6 | 6 | 1 |

b) ALL QTLs

| Pairwise Traits | nQTL1_L | nQTL2_L | nOverlap_L | Jaccard_Overlap |
| --- | --- | --- | --- | --- |
| CorollaLength-CorollaLength | 13 | 13 | 13 | 1 |
| CorollaLength-CorollaThroat | 13 | 14 | 8 | 0.42105263 |
| CorollaLength-CorollaWidth | 13 | 13 | 6 | 0.3 |
| CorollaLength-LongestStamenLength | 13 | 14 | 10 | 0.58823529 |
| CorollaLength-NectarVolume | 13 | 9 | 6 | 0.375 |
| CorollaLength-NectarySize | 13 | 16 | 7 | 0.31818182 |
| CorollaLength-OverallCorollaLength | 13 | 14 | 11 | 0.6875 |
| CorollaLength-PistilLength | 13 | 19 | 12 | 0.6 |
| CorollaLength-SeedLength | 13 | 7 | 4 | 0.25 |
| CorollaLength-SeedMass | 13 | 3 | 2 | 0.14285714 |
| CorollaLength-SeedWidth | 13 | 2 | 1 | 0.07142857 |
| CorollaLength-SepalLength | 13 | 11 | 9 | 0.6 |
| CorollaLength-SugarConcentration | 13 | 11 | 7 | 0.41176471 |
| CorollaThroat-CorollaThroat | 14 | 14 | 14 | 1 |
| CorollaThroat-CorollaWidth | 14 | 13 | 9 | 0.5 |
| CorollaThroat-LongestStamenLength | 14 | 14 | 7 | 0.33333333 |
| CorollaThroat-NectarVolume | 14 | 9 | 6 | 0.35294118 |
| CorollaThroat-NectarySize | 14 | 16 | 11 | 0.57894737 |
| CorollaThroat-OverallCorollaLength | 14 | 14 | 7 | 0.33333333 |
| CorollaThroat-PistilLength | 14 | 19 | 9 | 0.375 |
| CorollaThroat-SeedLength | 14 | 7 | 5 | 0.3125 |
| CorollaThroat-SeedMass | 14 | 3 | 3 | 0.21428571 |
| CorollaThroat-SeedWidth | 14 | 2 | 2 | 0.14285714 |
| CorollaThroat-SepalLength | 14 | 11 | 5 | 0.25 |
| CorollaThroat-SugarConcentration | 14 | 11 | 5 | 0.25 |
| CorollaWidth-CorollaWidth | 13 | 13 | 13 | 1 |
| CorollaWidth-LongestStamenLength | 13 | 14 | 7 | 0.35 |
| CorollaWidth-NectarVolume | 13 | 9 | 5 | 0.29411765 |
| CorollaWidth-NectarySize | 13 | 16 | 6 | 0.26086957 |
| CorollaWidth-OverallCorollaLength | 13 | 14 | 9 | 0.5 |
| CorollaWidth-PistilLength | 13 | 19 | 9 | 0.39130435 |
| CorollaWidth-SeedLength | 13 | 7 | 6 | 0.42857143 |
| CorollaWidth-SeedMass | 13 | 3 | 2 | 0.14285714 |
| CorollaWidth-SeedWidth | 13 | 2 | 1 | 0.07142857 |
| CorollaWidth-SepalLength | 13 | 11 | 6 | 0.33333333 |
| CorollaWidth-SugarConcentration | 13 | 11 | 3 | 0.14285714 |
| LongestStamenLength-LongestStamenLength | 14 | 14 | 14 | 1 |
| LongestStamenLength-NectarVolume | 14 | 9 | 8 | 0.53333333 |
| LongestStamenLength-NectarySize | 14 | 16 | 9 | 0.42857143 |
| LongestStamenLength-OverallCorollaLength | 14 | 14 | 11 | 0.64705882 |
| LongestStamenLength-PistilLength | 14 | 19 | 12 | 0.57142857 |
| LongestStamenLength-SeedLength | 14 | 7 | 7 | 0.5 |
| LongestStamenLength-SeedMass | 14 | 3 | 3 | 0.21428571 |
| LongestStamenLength-SeedWidth | 14 | 2 | 1 | 0.06666667 |
| LongestStamenLength-SepalLength | 14 | 11 | 7 | 0.38888889 |
| LongestStamenLength-SugarConcentration | 14 | 11 | 9 | 0.5625 |
| NectarVolume-NectarVolume | 9 | 9 | 9 | 1 |
| NectarVolume-NectarySize | 9 | 16 | 10 | 0.66666667 |
| NectarVolume-OverallCorollaLength | 9 | 14 | 7 | 0.4375 |
| NectarVolume-PistilLength | 9 | 19 | 7 | 0.33333333 |
| NectarVolume-SeedLength | 9 | 7 | 3 | 0.23076923 |
| NectarVolume-SeedMass | 9 | 3 | 3 | 0.33333333 |
| NectarVolume-SeedWidth | 9 | 2 | 1 | 0.1 |
| NectarVolume-SepalLength | 9 | 11 | 5 | 0.33333333 |
| NectarVolume-SugarConcentration | 9 | 11 | 5 | 0.33333333 |
| NectarySize-NectarySize | 16 | 16 | 16 | 1 |
| NectarySize-OverallCorollaLength | 16 | 14 | 5 | 0.2 |
| NectarySize-PistilLength | 16 | 19 | 9 | 0.34615385 |
| NectarySize-SeedLength | 16 | 7 | 7 | 0.4375 |
| NectarySize-SeedMass | 16 | 3 | 5 | 0.35714286 |
| NectarySize-SeedWidth | 16 | 2 | 2 | 0.125 |
| NectarySize-SepalLength | 16 | 11 | 7 | 0.35 |
| NectarySize-SugarConcentration | 16 | 11 | 9 | 0.5 |
| OverallCorollaLength-OverallCorollaLength | 14 | 14 | 14 | 1 |
| OverallCorollaLength-PistilLength | 14 | 19 | 12 | 0.57142857 |
| OverallCorollaLength-SeedLength | 14 | 7 | 5 | 0.3125 |
| OverallCorollaLength-SeedMass | 14 | 3 | 2 | 0.13333333 |
| OverallCorollaLength-SeedWidth | 14 | 2 | 1 | 0.06666667 |
| OverallCorollaLength-SepalLength | 14 | 11 | 8 | 0.47058824 |
| OverallCorollaLength-SugarConcentration | 14 | 11 | 7 | 0.38888889 |
| PistilLength-PistilLength | 19 | 19 | 19 | 1 |
| PistilLength-SeedLength | 19 | 7 | 7 | 0.36842105 |
| PistilLength-SeedMass | 19 | 3 | 3 | 0.15789474 |
| PistilLength-SeedWidth | 19 | 2 | 1 | 0.05 |
| PistilLength-SepalLength | 19 | 11 | 9 | 0.42857143 |
| PistilLength-SugarConcentration | 19 | 11 | 8 | 0.36363636 |
| SeedLength-SeedLength | 7 | 7 | 7 | 1 |
| SeedLength-SeedMass | 7 | 3 | 3 | 0.42857143 |
| SeedLength-SeedWidth | 7 | 2 | 2 | 0.28571429 |
| SeedLength-SepalLength | 7 | 11 | 6 | 0.5 |
| SeedLength-SugarConcentration | 7 | 11 | 4 | 0.28571429 |
| SeedMass-SeedMass | 3 | 3 | 3 | 1 |
| SeedMass-SeedWidth | 3 | 2 | 2 | 0.66666667 |
| SeedMass-SepalLength | 3 | 11 | 2 | 0.16666667 |
| SeedMass-SugarConcentration | 3 | 11 | 1 | 0.07692308 |
| SeedWidth-SeedWidth | 2 | 2 | 2 | 1 |
| SeedWidth-SepalLength | 2 | 11 | 2 | 0.18181818 |
| SeedWidth-SugarConcentration | 2 | 11 | 0 | 0 |
| SepalLength-SepalLength | 11 | 11 | 11 | 1 |
| SepalLength-SugarConcentration | 11 | 11 | 5 | 0.29411765 |
| SugarConcentration-SugarConcentration | 11 | 11 | 11 | 1 |

**Table S6** Average predicted genetic correlations, *r_Q_*, within and between modules with standard errors (in parentheses). a) Average predicted *r_Q_* from conservative GWS-only QTLs. b) Average predicted *r_Q_* from ALL QTLs.

a) GWS QTLs

MODULE       flower                  nectar                      seed

    flower      0.4692 (0.0454)     0.1856 (0.0192)    -0.1899 (0.0368)

    nectar      0.1856 (0.0192)     0.1789 (0.1220)    -0.0968 (0.0539)

    seed	     -0.1899 (0.0368)    -0.0968 (0.0539)     0.1870 (0.1870)

b) ALL QTLs

MODULE          flower                 nectar                      seed

    flower      0.5537 (0.0419)     0.3056 (0.0296)    -0.3472 (0.0276)

    nectar     0.3056 (0.0296)     0.3104 (0.1217)    -0.3242 (0.0668)

    seed    -0.3472 (0.0276)    -0.3242 (0.0668)     0.6602 (0.1009)

**Table S7** Randomization test (random placement of QTLs in genome) of whether QTLs within modules are spatially clustered in the genome. 1,000 permutations were run and the average overlap within modules calculated for each permutation. a) GWS (genome-wide significant) QTLs. b) ALL QTLs.

a) GWS QTLs

Overall test: P < 0.0001

Flower module: P < 0.0001

Nectar module: P = 0.355

Seed module: P = 0.175

b) ALL QTLs

Overall test: P < 0.0001

Flower module: P < 0.0001

Nectar module: P < 0.032

Seed module: P < 0.002

**Table S8** Number of QTLs with effects in same direction as or opposite direction from difference between species. Proportions are not significantly different (Fisher exact test, P = 0.1114).

QTL Type	Same direction	Opposite direction	Proportion in same direction

GSW	76	22	0.776

CSW	43	5	0.895

GWS + CWS	119	27	0.815

**Table S9** Contra-directional QTLs summary. Numbers and proportions of floral and nectar QTLs with effects in the direction opposite species difference that overlap with at least one other floral or nectar QTL with effects in the same direction as species difference. Calculations based on ALL QTLs.

| **Trait** | **Total number of QTLs** | **Number of Contra QTLs** | **Number of Contra QTLs overlapping consistent floral QTLs** | **Number of Contra QTLs overlapping consistent nectar QTLs** | **Number of Contra QTLs overlapping either** |
| --- | --- | --- | --- | --- | --- |
| Corolla Width | 13 | 0 | NA | NA | NA |
| Corolla Throat | 14 | 3 | 2 | 1 | 2 |
| Corolla Length | 13 | 2 | 1 | 2 | 2 |
| Overall Corolla  Length | 14 | 1 | 1 | 1 | 1 |
| Sepal Length | 11 | 5 | 3 | 3 | 4 |
| Longest Stamen  Length | 14 | 2 | 1 | 2 | 2 |
| Pistil Length | 19 | 4 | 0 | 3 | 3 |
| Nectar Volume | 9 | 1 | 0 | 1 | 1 |
| Nectar Sugar  Concentration | 11 | 1 | 0 | 1 | 1 |
| Nectary Size | 16 | 4 | 0 | 1 | 1 |
|  | TOTAL | 23 | 8 | 15 | 17 |
|  | Proportion of total | | 0.3478 | 0.6522 | 0.7391 |
